# Cost-function Optimized Maximal Overlap Drift Estimation for Single Molecule Localization Microscopy

**DOI:** 10.64898/2026.03.27.714864

**Authors:** Lenny Reinkensmeier, Sarah Aufmkolk, Irene Farabella, Alexander Egner, Mark Bates

## Abstract

Single-molecule localization microscopy (SMLM) methods enable fluorescence imaging of biological specimens with nanometer-scale resolution. Although fluorophore localization precision is theoretically limited only by photon statistics, in practice the resolution of SMLM images is often degraded by physical drift of the sample and/or the microscope during data acquisition. At present, correcting this effect requires either specialized stabilization systems or computationally intensive post-processing, and established drift correction algorithms based on image cross-correlation suffer from limited temporal resolution. In this study we introduce COMET, a new method for SMLM drift estimation which achieves a substantially higher precision, accuracy, and temporal resolution compared with existing algorithmic approaches. We demonstrate that improved drift estimation translates directly into higher SMLM image resolution, limited by localization precision rather than drift artifacts. COMET is applicable to all types of SMLM data, operating directly on 2D or 3D localization datasets, and is readily integrated into analysis workflows. We benchmark its performance using both simulations and experiments, including STORM, MINFLUX, and Sequential OligoSTORM measurements, where long acquisition times make drift correction particularly challenging. COMET is published as an open-source, Python-based software project and is also available on open cloud-computing platforms.

---

Fluorescence imaging techniques based on single molecule localization microscopy (SMLM) have pushed the limits of image resolution in biological specimens to sub-10 nanometer length-scales and beyond.^1-4^ The underlying concept of SMLM is to measure the spatial coordinates of sparse subsets of fluorophores at different time points, gradually accumulating molecular localizations which form an image. During the measurement, however, mechanical drift of the sample stage, mirrors, or other changes in the optical paths cause unavoidable shifts in the position of the sample with respect to the microscope, cumulatively spreading out the localizations in time and thereby systematically distorting the reconstructed image.

Several approaches have been introduced to minimize and correct for drift, including innovative microscope design,^5^ fiducial marker tracking,^4, 6, 7^ and cross-correlation of localization data across multiple time segments^8-10^. However, each of these approaches presents limitations in terms of practicality, accuracy, and computational efficiency. Novel microscope designs have proven beneficial but are insufficient to fully eliminate drift at the nanometer scale.^5^ Fiducial marker tracking, while highly accurate, is challenging in many imaging contexts, for example due to the requirement for the marker to be in the same focal plane as the sample.^4^ Correlation-based post-processing approaches are more generally applicable because they rely only on the localization coordinates. In these methods, the localization dataset is divided into successive time segments and spatial shifts between segments are estimated using image cross-correlation.^8-10^ However, statistically robust correlations require a sufficient number of localizations in each segment, resulting in a drift estimate that is relatively coarse-grained in time. As a result, such approaches are unable to correct for fast drift transients.

In this work, we develop a new approach to drift correction of SMLM data, which requires no fiducial markers and yet enables accurate drift estimation with high temporal resolution. Our method, Cost-function Optimized Maximal overlap drift EsTimation (COMET), uses numerical optimization to determine the unique time-varying displacement vector that maximizes the spatial overlap of the localization coordinates. Testing confirmed that this vector corresponds to the true experimental sample drift, and we found that the accuracy and speed of the COMET algorithm strongly outperforms currently established drift correction methods. We present the principles of COMET, validate its performance using simulated data and real-data benchmarking, and demonstrate its use in nanoscale bio-imaging applications.

## Results

### Principle and optimization framework of COMET

To overcome the drawbacks of fiducial marker tracking and cross-correlation-based drift correction, we hypothesized that a more efficient approach would operate directly on the localization coordinates while avoiding the necessity to generate intermediate SMLM images. Our method, COMET, is based on two postulates: *(i)* A unique, time-varying displacement vector 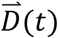 exists, which, when subtracted from the localization coordinates, shifts the localizations such that they maximally overlap in space; *(ii)* Assuming a sufficient number of localizations at each time point, this 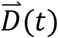 corresponds to the true sample drift during the experiment. Thus, we hypothesize that the sample drift can be estimated by solving for the displacement vector that maximizes the spatial overlap of the localizations.

To determine the displacement vector 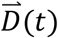, we formulated an optimization problem based on a cost function (Equation 1) that quantitatively describes the spatial overlap of the localizations, modeled as Gaussian functions (Fig. 1a). Given the measured localization coordinates 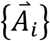, the COMET cost function *f*_*c*_ and its derivatives are given by

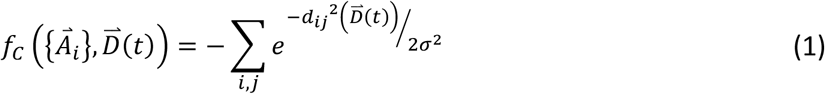

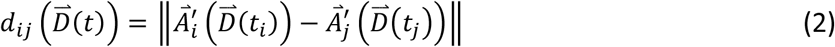

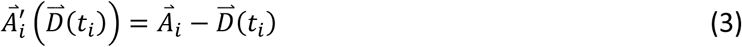

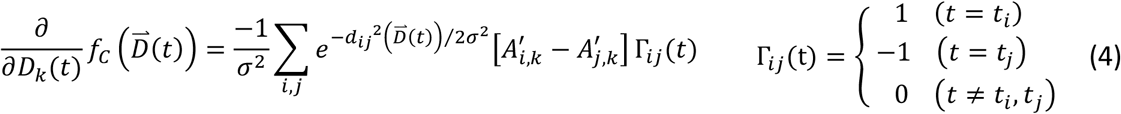

where *i* and *j* are localization indices, *k* is a spatial dimension index, *d_ij_* is the distance between localizations *i* and *j* (Equation 2), 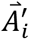 are the localization coordinates after subtraction of the drift (Equation 3), *t_i_* are the time coordinates of the localizations, and *σ* is a Gaussian length-scale that defines the effective interaction distance between pairs of localizations.

**Figure 1:**
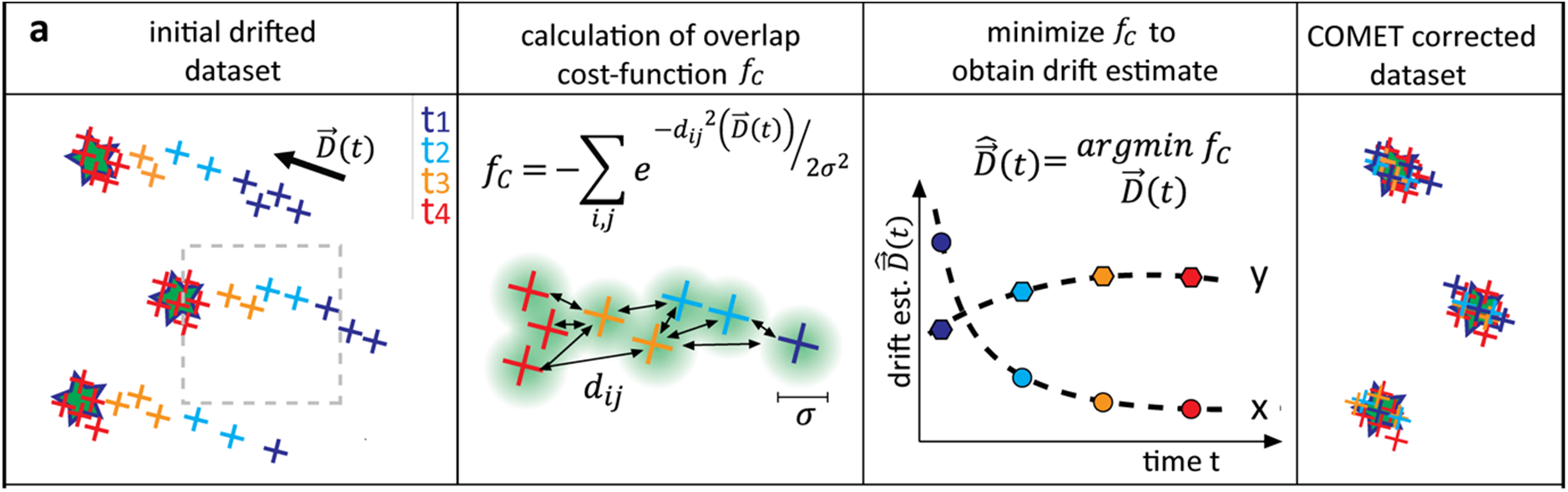
COMET principle. **(a)** Illustration of three emitters (stars) affected by drift. The localizations of each emitter across fourtimepoints (t1-t4) are shown as color-coded crosses with a 2D Gaussian representing uncertainty, spread out due to the drift. COMET identifies pairs of localizations from different timepoints and optimizes for the drift estimate vector 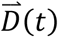 that maximizes their spatial overlap and effectively drift corrects the dataset.

The calculation of *f*_*c*_ presents computational challenges due to the number of terms in the sum over coordinate pairs, which may reach 10^12^ or more for a typical SMLM measurement. However, pairs separated by distances much larger than *σ* contribute negligibly to the result and can be safely excluded. We therefore define a maximum expected drift *D*_*max*_ and exclude coordinate pairs whose initial separations exceed this distance. Despite this simplification, for typical datasets the number of pairs may still reach orders of 10^8^. To keep the computation feasible, we parallelized the computation of *f*_*c*_ and its derivatives (Equation 4) using GPU-based processing, leveraging the fact that the terms of the sums in Equations 1 and 4 are independent and separable.

The drift estimate 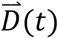 is then obtained by minimizing *f*_*c*_ with respect to the drift trajectory. To efficiently manage the high dimensionality of the parameter space, we employed the limited-memory Broyden–Fletcher–Goldfarb–Shanno algorithm (L-BFGS-B), which is well-suited for large-scale optimization problems.^11^ To facilitate convergence to the global minimum, we developed a protocol in which the Gaussian scale parameter *σ* is iteratively reduced over multiple rounds of optimization. Beginning with a relatively large value for *σ* (approximately 1/3 of *D*_*max*_), the minimizer initially converges to the vicinity of the global minimum by correcting for the large length-scale features of the drift trajectory. As the value of *σ* is reduced, the cost function *f*_*c*_ becomes more sensitive to drift on shorter length-scales, allowing the optimizer to converge to a progressively more accurate solution. The procedure is repeated until a stopping criterion is reached, when no further improvement in the drift estimate is observed (see full workflow in Supplementary Information, Supplementary Fig. 1).

### Estimation of sample drift in experimental SMLM data

To assess the performance of COMET on experimental data, we applied the algorithm to a 3D SMLM dataset of nuclear pore complexes (NPCs) acquired using 4Pi-STORM (Fig. 2a).^2^ We compared the resulting drift estimate with that obtained using redundant cross-correlation (RCC),^10^ a widely used correlation-based drift correction method.

**Figure 2:**
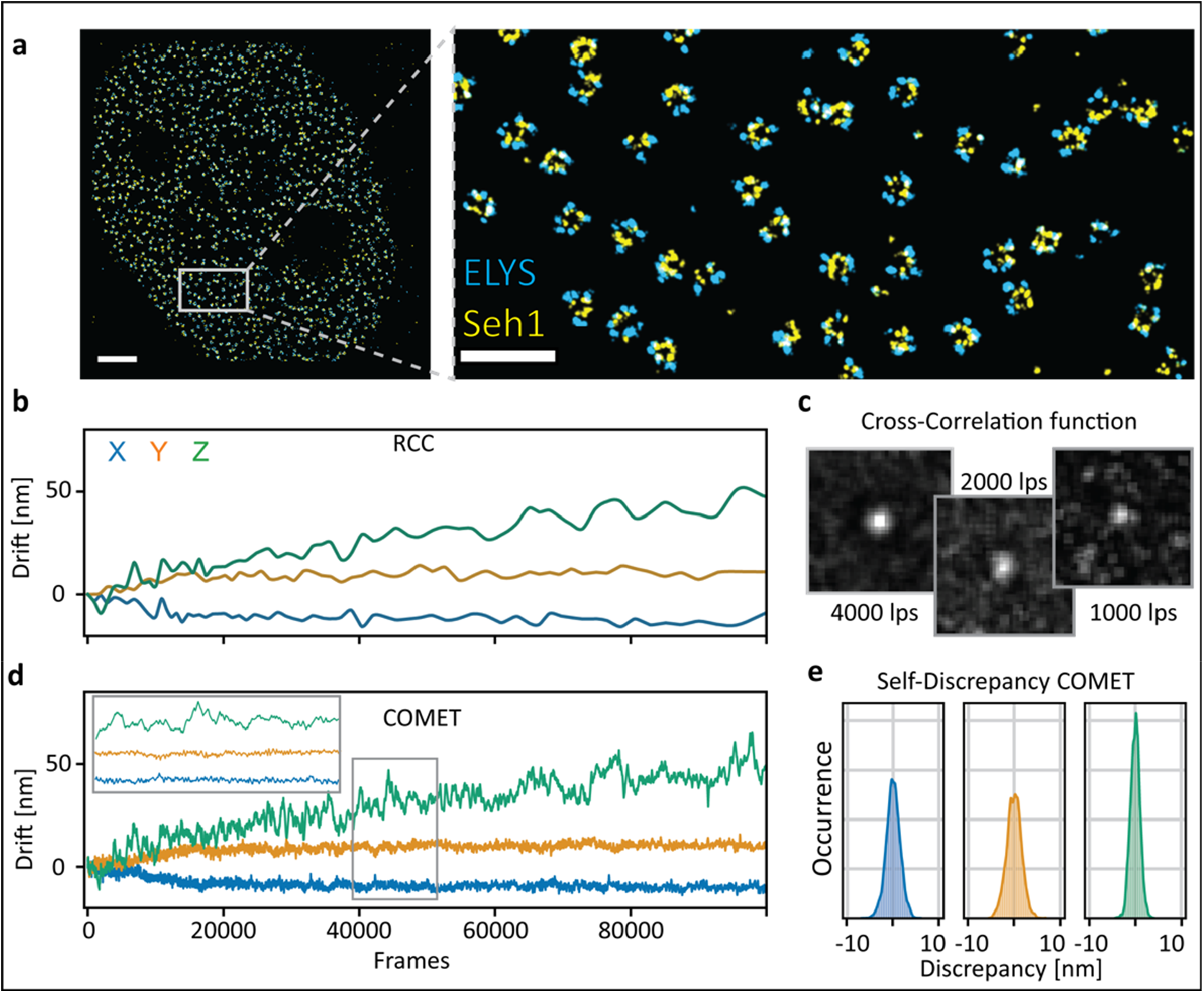
Comparison of COMET and RCC drift correction on 4Pi-STORM data. **(a)** Rendering of the dataset showing NPCs in the nuclear membrane of a U-2-OS cell. The nucleoproteins ELYS and Seh1 are shown in cyan and yellow, respectively. **(b)** Estimated drift vs. frame number found for RCC. **(c)** 2D-Projection renderings of cross-correlation function centers of an example segment pair for the RCC drift correction of the presented dataset for three different segmentation conditions using 4000, 2000, and 1000 localizations per segment (lps). 4000 localizations per segment corresponds to the segmentation used to retrieve (b). **(d)** Estimated drift vs. frame number found for COMET with the inset showing a close-up of the marked region (grey square) to highlight the fine temporal details. **(e)** Self-consistency check performed for the COMET drift estimate using the same number of localizations per segment as in (d), 50. For this, two independent halves of the dataset were corrected independently, and a histogram of the resulting discrepancy in the drift estimates is shown for X, Y, and Z, respectively. Scalebars in (a) 2 µm and 200 nm for overview and inset, respectively.

The NPC dataset was first processed using RCC with a time segment size of 4000 localizations per segment, chosen to represent a stable and commonly used operating regime for correlation-based drift correction. This yielded 68 time segments, requiring a total of 2278 three-dimensional cross-correlations and resulting in a computation time of approximately 8 minutes. The resulting drift estimate (Fig. 2b) reveals a slow drift of the sample in the x, y, and z dimensions, with superimposed fluctuations with an amplitude of ∼10 nm. Reducing the segment size further degraded the stability of the RCC solution; below approximately 2000 localizations per segment the drift trajectories became unstable and produced spurious outliers due to excessive noise in the correlation function (Fig 2c; Supplementary Fig. 2).

We then applied COMET to the same dataset, selecting a substantially smaller time segment size of 250 localizations per segment, corresponding to approximately 1100 time segments, using *D*_*max*_ = 300 nm, and an initial Gaussian scale parameter *σ* = 30 nm. Starting from an initial drift estimate of 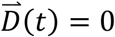, the optimizer converged rapidly to a stable result. Successive optimization rounds with progressively reduced Gaussian scale parameter *σ* consistently refined the solution until a final drift estimate was reached after a total computation time of approximately 18 seconds. The COMET drift estimate (Fig. 2d, Supplementary Fig. 3) exhibits markedly higher temporal resolution and clearly resolves fast nanoscale drift transients that are fully obscured in the RCC solution.

Although the COMET solution is consistent with the low-resolution RCC result, we asked whether the fast transients detected by COMET were real or artefactual. To assess this, we divided the NPC dataset into two independent parts (localizations first detected in even versus odd frames of the raw data), estimated the drift independently for each subset using COMET, and compared the resulting drift trajectories. The discrepancy between the independent solutions was on the order of 1 nm (Fig. 2e; Supplementary Fig. 4), demonstrating both the precision of the COMET estimate and the reliability of the resolved fast drift transients.

Finally, we asked how far the temporal resolution of the drift estimate could be pushed. Using the self-consistency test as a quality metric, we increased the number of time segments in the solution and explored the practical performance limits of RCC in parallel. Using this approach, we found that the temporal resolution of RCC could be pushed to approximately 2000 localizations per segment (136 segments, >3 hours computation time), beyond which the RCC solution diverged. In contrast, COMET continued to converge at time resolutions as fine as 50 localizations per segment while maintaining a drift uncertainty on the order of 1 nm.

Remarkably, the total computation time of COMET remained essentially unchanged (≈18 seconds) as the temporal resolution increased. Overall, this corresponds to an improvement in temporal resolution of approximately forty-fold relative to the best-performing RCC solution, while the total computation time of COMET was approximately 500-fold faster.

### Estimation of simulated drift

To validate the accuracy of COMET under conditions where the true sample drift is known, we evaluated the algorithm using simulated SMLM datasets containing artificially induced drift. Unlike experimental data, simulated datasets provide direct access to the ground truth solution, allowing the accuracy of the recovered drift to be quantified explicitly. To generate realistic test data, we started from a real STORM image of microtubules obtained from the Shareloc open SMLM image server (https://shareloc.xyz),^12^ which was randomly resampled to generate a localization dataset representative of an experimentally observed structure (Fig. 3a,b; see Supplementary Information; Supplementary Fig 5). A three-dimensional drift trajectory was generated and applied to the data, incorporating both slow and fast oscillatory components to approximate experimentally observed drift dynamics across multiple timescales (Supplementary Fig. 6).

**Figure 3:**
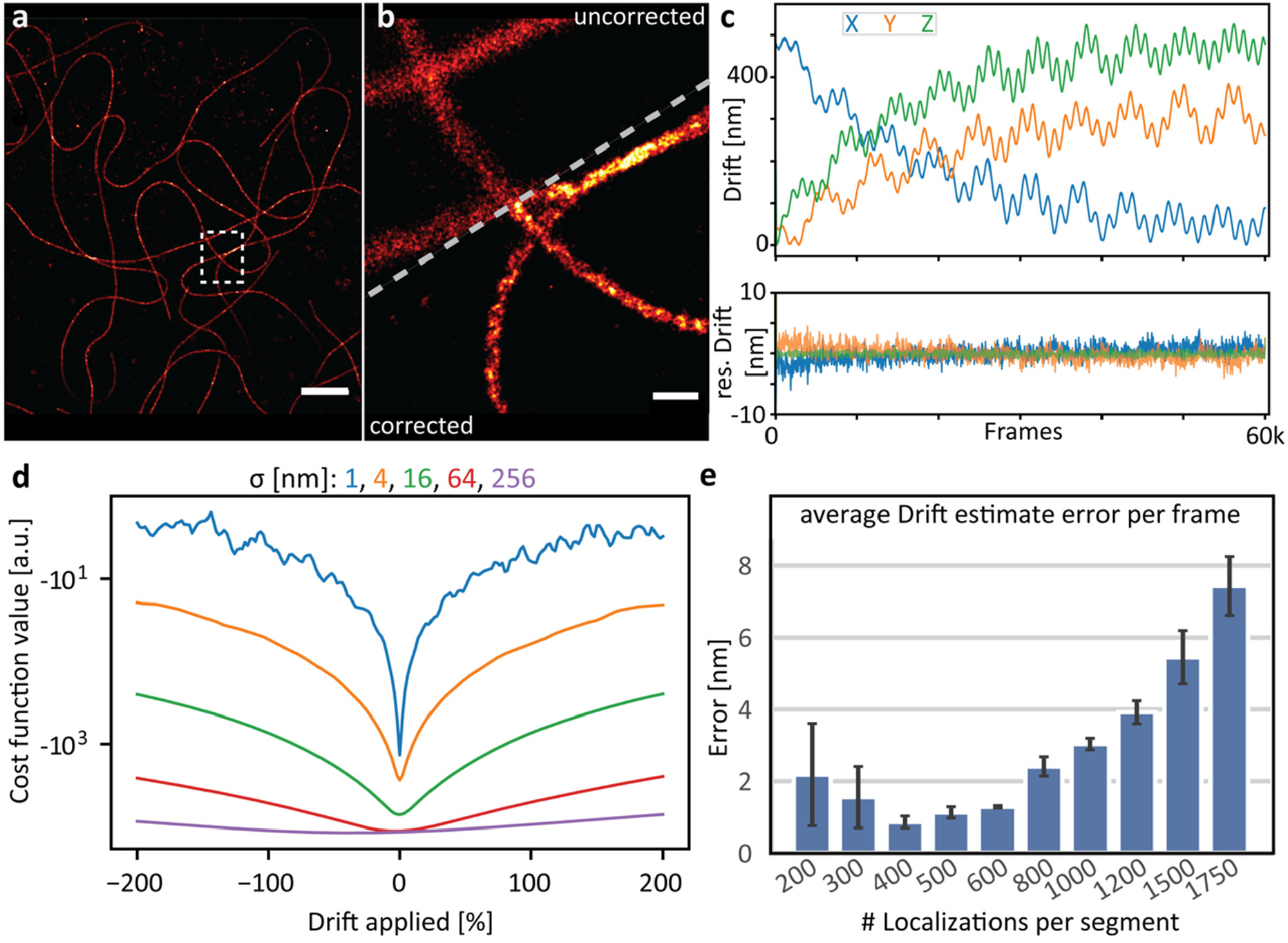
COMET proof-of-concept demonstration. **(a)** Rendering of resampled STORM dataset showing tubulin filaments. **(b)** Rendered close-up of the region marked by the white square in (a) before and after the COMET drift correction was applied. **(c)** Aritificially generated drift curves, comprising a fast and slow oscillation, which was applied to the resampled STORM dataset shown in (a, b). **(d)** The cost function landscape as a function of applied drift curves shown in (c) for different Gaussian length scale parameters. **(e)** The error of the COMET drift estimate as function of segmentation of the dataset shown in (a). Panel Scalebars in (a) and (b) are 2 µm and 250 nm respectively.

After adding the simulated drift to the data, the localization list was processed using COMET with a time segment size of 200 localizations per segment, a maximum expected drift *D*_*max*_ = 600 nm, and an initial Gaussian scale parameter *σ* = 200 nm, following the optimization strategy established on experimental data. The recovered drift estimate was compared directly with the ground-truth drift. The residual error of the COMET drift estimate was approximately 2 nm or lower (Fig. 3c). Notably, the residual error in the *x* and *y* dimensions exceeded that in *z*. Since the dataset is effectively two-dimensional, residual errors in x and y are expected to be limited by the intrinsic localization precision of the data, whereas the residual error in z likely reflects the precision limit of the COMET algorithm under these conditions.

To better understand the behavior of the COMET optimization and the role of the Gaussian scale parameter *σ*, we investigated the high-dimensional landscape of the COMET cost function. We applied scaled versions of the simulated drift to the microtubule dataset and computed the corresponding values of *f*_*c*_. By varying the scaling factor between −2 and +2, where +1 corresponds to the full simulated drift and 0 to no drift, we visualized a one-dimensional slice of the cost-function landscape between the initial condition and the ground-truth solution. This calculation was repeated for different values of *σ* (Fig. 3d; Supplementary Fig. 7a,b). These results show that for larger values of *σ*, the landscape becomes smoother, facilitating convergence toward the global minimum. As *σ* is reduced, the cost function becomes more sensitive to short length-scale drift variations, allowing the optimizer to refine the solution with higher precision. Importantly, the minimum of *f*_*c*_ occurs at the zero-drift condition, confirming an underlying assumption of COMET.

We next investigated how the temporal resolution of the drift estimate depends on the time step size. The information content of each time segment depends on the spatial structure of the sample, and the optimal time step size is therefore sample dependent. Point-like features require fewer localizations to determine their spatial position, whereas continuous or filamentous structures with diffuse boundaries require higher localization densities per time segment. Using the simulated microtubule dataset, which represents filamentous structures, we evaluated the residual error in the COMET drift estimate as a function of time step size. As the step size decreased, the residual error initially improved due to higher temporal resolution, but increased again once the number of localizations per segment became insufficient (Fig. 3e). For this dataset, the optimal performance was obtained at approximately 400 localizations per segment. As a practical guideline, a few hundred localizations per segment provide robust performance for most SMLM datasets tested.

### Benchmark comparison with existing drift correction methods

Following its strong performance relative to RCC, we benchmarked COMET against a broader set of published SMLM drift correction methods. The comparison included the recently introduced AIM^13^ and Minimum Entropy (ME)^14^ approaches, as well as correlation-based methods including redundant cross-correlation (RCC) and direct cross-correlation (DCC),^8, 9^ in which spatial shifts are estimated sequentially between adjacent time segments. For the correlation-based methods, we tested both the RCC implementation available in the SMAP analysis package^15^ and our own optimized implementations of RCC and DCC.

To account for differences in structural content and localization density, we generated three simulated 3D SMLM datasets representing densely, moderately, and sparsely sampled structures (Fig. 4a; see Supplementary Information). For each method and dataset, a range of time segment sizes was tested, and the best-performing result was identified by minimizing the mean distance between drift-corrected localizations and their ground-truth positions (Supplementary Tables 1-3)). From this analysis, we extracted two metrics for each method: the precision of the optimal solution and the corresponding computation time (Fig. 4b,c,d; Supplementary Fig. 9).

**Figure 4:**
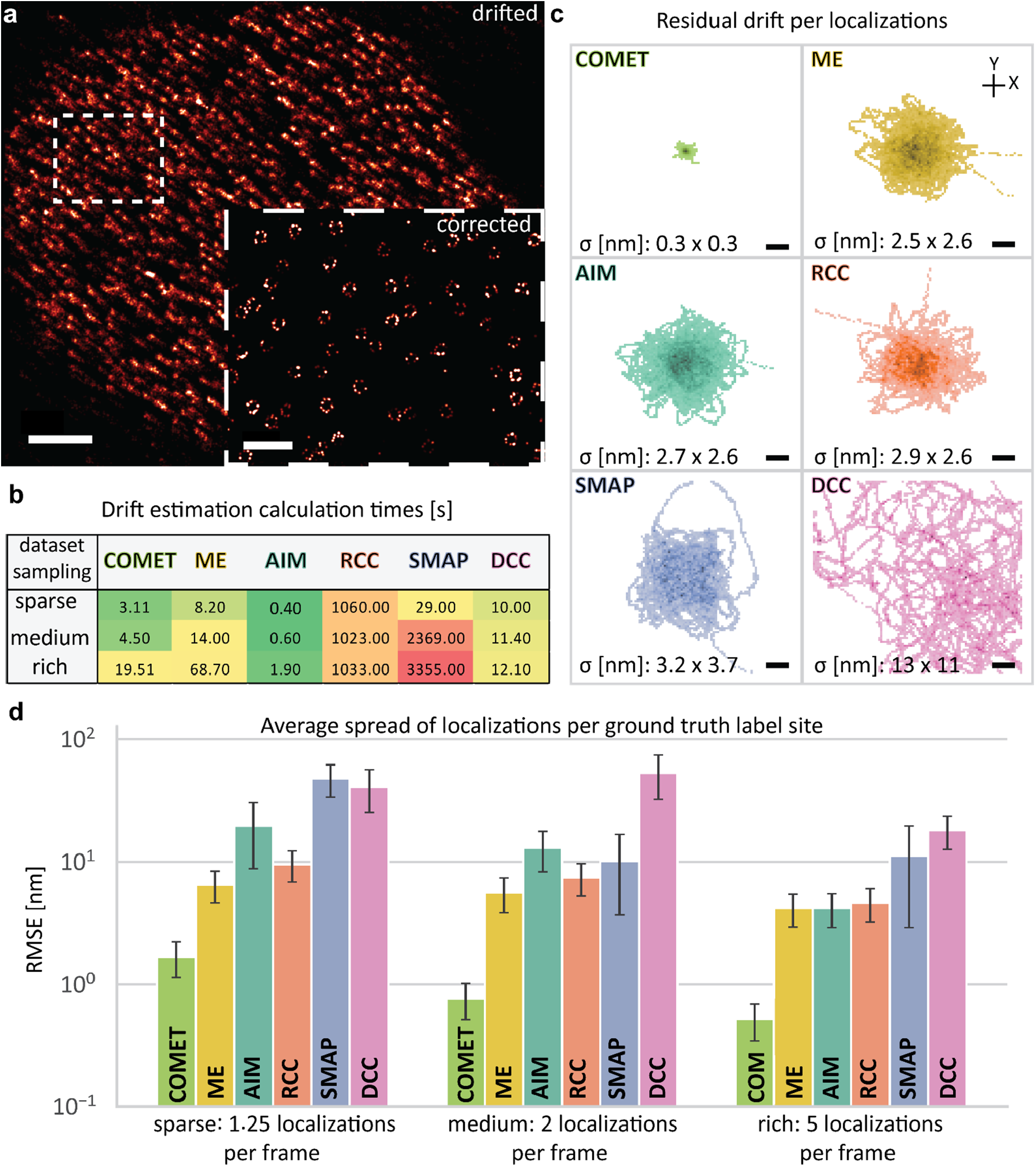
Benchmarking COMET against established methods. To quantitatively assess the drift correction performance, a real 3D NPC dataset recorded with 4Pi-STORM was resampled under three conditions, varying the number of localizations in the dataset. The three simulated datasets, called *sparse, medium*, and *rich*, corresponded to low, medium, and high densities of localizations per unit time. Artificial drift was applied to the datasets, and the drift was then estimated using each of the tested methods: COMET, AIM, Minimum Entropy (ME), redundant cross-correlation (RCC), and direct cross-correlation (DCC). **(a)** Image of the resampled dataset, showing the uncorrected and drift-corrected data. Scalebars: 2 µm in overview, 500 nm in inset. (**b)** Computation time in seconds for the best-case correction of each method. Color indicates calculation performance from green (fastest) to red (slowest). **(c)** Each method was tested using temporal segmentation of 4 to 960 localizations per time segment and the best-performing result per (i.e., lowest drift discrepancy relative to the ground truth) was selected. The color-coded 2D-Histograms show the residual drift per localization best-case result of each method for the rich sampling scenario. **(d)** Histogram showing the best-case drift correction performance per method and sampling condition, evaluated as the average spread of localizations to the respective ground truth label-site due to drift. Scalebars in (c) 1 nm.

In terms of drift estimation precision, COMET consistently outperformed all other methods by a factor of approximately 4–50 across all simulated conditions. The Minimum Entropy method yielded the next best performance, consistent with its conceptually related formulation. In contrast, correlation-based approaches (RCC, SMAP-RCC, and DCC) showed substantially lower precision, in agreement with the limitations identified above.

When comparing computation times, AIM was the fastest method overall, but its precision was an order of magnitude worse than that of COMET in all cases. DCC also performed relatively quickly, reflecting the simplicity of the algorithm, but exhibited the poorest precision among the tested methods. COMET and ME showed comparable computation times, typically ranging from a few seconds to approximately one minute. Finally, the unfavorable scaling of RCC with increasing temporal resolution was evident, with both SMAP-RCC and our optimized RCC implementation requiring more than one hour to process the medium and large simulated datasets. Together, these results demonstrate that COMET provides both the highest drift estimation precision and competitive computation times across a wide range of imaging conditions.

### COMET drift correction of Sequential OligoSTORM data

OligoSTORM and OligoDNA-PAINT are super-resolution imaging techniques for mapping the three-dimensional organization of chromosomes within the nucleus using Oligopaint FISH probes labeled with photo-switchable fluorophores or short DNA docking strands.^16-18^ These experiments, when coupled with multiple rounds of acquisitions, typically involve imaging large nuclear volumes over extended acquisition times, including repeated axial scanning and multiple rounds of hybridization via readout-probe exchange.^19, 20^ As a result, sample drift introduced by focal shifts and buffer exchanges presents a significant challenge and must be accurately corrected to enable reliable reconstruction of chromosome territories.

Previous Sequential OligoSTORM studies have relied on fiducial marker–based drift correction.^18^ However, this approach requires manual identification and tracking of suitable fiducials and may fail when fiducial markers move out of focus during axial scanning. To assess whether COMET can provide a robust, fiducial-free alternative for large genomic SMLM datasets, we applied the algorithm to sequential OligoSTORM data acquired over three hybridization rounds targeting the same genomic region (chr2:112.67–113.17 Mb in human RPE-1 nuclei) (Fig. 5a, each round represented by a different color). Each round comprised repeated axial scans spanning approximately 4 μm, resulting in extended acquisition times spanning over 500.000 camera frames.

**Figure 5:**
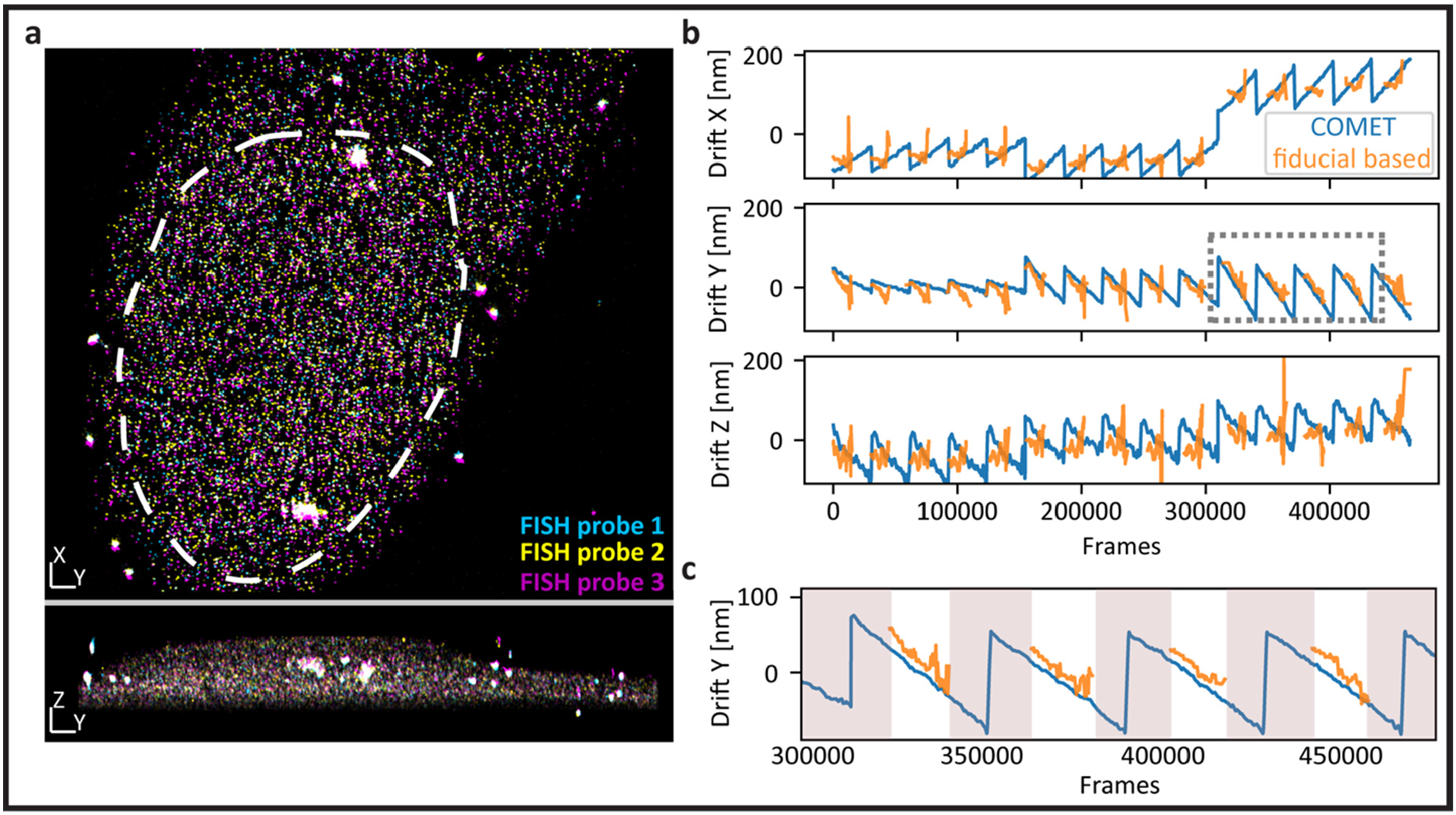
Results of COMET correction on OligoSTORM dataset in comparison to fiducial marker-based drift correction. **(a)** Sequential OligoSTORM image, before drift correction, showing a cell nucleus labeled with Oligopaint probes against a 500kb region on chromosome 2 (chr2:112.67–113.17 Mb) in human RPE-1 nuclei. Three rounds of targeting the same segment are shown in different colors, and the outline of the nucleus is marked with a dashed line. The upper panel shows the x-y view, and the lower panel shows the y-z profile view. **(b)** Drift estimate vs. frame number for both methods. **(c)** Close up of the region in (b) marked with a dashed grey square. Time segments in which the fiducial-based method lost track are highlighted in grey. Lengths of axis markers in (a) 1 um.

The Sequential OligoSTORM data from each hybridization round were corrected independently using COMET with a fixed time segment size of 250 frames, corresponding to approximately 600 localizations per segment. The resulting drift trajectories are shown in Fig. 5b, together with drift estimates obtained using fiducial marker tracking computed by the Bruker SRX software (see Supplemental Methods). A prominent feature of the COMET drift estimate is the periodic displacement induced by axial focal plane scanning. This oscillatory drift is largely absent from the fiducial-based correction, because fiducial markers are intermittently lost when they move out of focus during the scan (Fig. 5c). Additionally, the fiducial-based correction exhibits significant noise spikes, which may arise due to beads which are not stably bound to the sample.

To quantify the impact of drift correction, in terms of localization accuracy, we compared the localization coordinates obtained after COMET correction with those obtained using fiducial-based correction. The resulting discrepancies are shown in Supplemental Fig. 10. Across all spatial dimensions, a substantial fraction of localizations differed by 50 nm or more between the two correction methods. Together, these results highlight the ability of COMET to provide reliable drift correction in large and extended SMLM datasets, featuring large-amplitude rapidly varying drift patterns, such as those encountered in genomic super-resolution imaging. Furthermore, in this example COMET outperformed the fiducial-based method, showcasing the robustness of the COMET approach.

## Discussion

Since the early conception of SMLM methods, correcting sample drift has been a persistent challenge. Streaks of localizations in STORM, PALM, DNA-PAINT, and MINFLUX images starkly illustrate how even subtle mechanical or thermal instabilities can degrade image quality at nanometer length scales, regardless of the sophistication of the microscope design. Approaches based on fiducial marker tracking and image correlation have provided practical and robust solutions. However, as SMLM techniques now routinely reach single-nanometer localization precision and beyond,^3,21^ drift correction methods capable of capturing rapid drift transients with correspondingly high spatial and temporal resolution have become essential.

The advantages of COMET arise from its efficient use of localization data. By directly optimizing a cost function defined on localization coordinates, COMET avoids intermediate image rendering and makes more effective use of the available information than correlation-based approaches. As demonstrated here, this leads to substantially improved temporal resolution of the drift estimate, enabling accurate correction of fast drift dynamics while simultaneously reducing computation time. In practice, these gains translate directly into improved effective SMLM resolution, with image quality limited primarily by localization precision rather than residual drift artifacts.

In this work, we introduced the COMET algorithm and validated its performance using both simulated and experimental datasets, including systematic benchmarking against established and recently developed drift correction methods. Across all tested conditions, COMET consistently achieved the most accurate drift estimates, resulting in the highest reconstructed image quality while remaining computationally efficient even when high-accuracy results were required. Robust convergence of the optimization was observed across a wide range of datasets, facilitated by iterative refinement of the Gaussian length-scale, which adapts naturally to varying localization densities. This performance may open new possibilities for advanced drift-correction modalities, for example, in dynamic (live-cell) SMLM data or in field-dependent drift estimation of optically aberrated images.

To facilitate adoption of the method, we have developed a Python-based implementation of COMET that can be simply integrated into analysis workflows. The software is publicly available and has already been incorporated into the widely used Picasso analysis framework.^22^ Beyond the datasets presented here, COMET has been successfully applied to diverse SMLM modalities including MINFLUX and 4Pi-STORM, as well as to data acquired under extreme drift conditions (Supplementary Figs. 11-13), confirming its broad applicability. To enable use without direct access to GPUs, a freely accessible COMET server has been established at https://comet.smlm.tools (Supplementary Fig. 14).

