## Supplementary Information for "Cost-function Optimized Maximal Overlap Drift Estimation for Single Molecule Localization Microscopy"

### Methods

#### Supplementary Implementation Details

##### COMET core implementation

The presented COMET implementation calculates a cost function representing the spatial overlap of localization pairs, optimized using the L-BFGS-B algorithm. The overlap is quantified by a Gaussian mixture model based on a single parameter, termed the Gaussian length-scale $\sigma$, which modulates the effective overlap as a function of the physical distances between each pair of localizations. Iterative minimization of the cost function for decreasing Gaussian scale parameter is employed to refine the drift estimate until convergence (typical initial scale 100 nm or one third of the expected drift, target scale <10 nm). After a successful minimization step the interaction length parameter is reduced by a factor of 1.5. Convergence is defined as the point at which further reduction of the Gaussian scale parameter no longer refines the drift estimate. Accordingly, convergence is reached when the change in the drift estimate between successive minimization steps decreases below this criterion, with the final step satisfying this condition. if the L-BFGS-B optimizer fails, the minimization step is (i) repeated with the Gaussian length-scale increased by a factor of two after two overall failures or (ii) the drift estimation is terminated after the fifth failure.

##### COMET workflow

Details of the workflow are shown in supplementary Fig. 1. In the first two steps the raw single molecule localization microscopy (SMLM) dataset is segmented into time windows and localization pairs within a range of the maximum expected drift are identified. The coordinates and pairs are then loaded into GPU memory and the core of the COMET algorithm estimates the drift vector that maximizes the overlap iteratively refined at length scales in a nested loop as described above. Optionally, the drift estimate is smoothed using boxcar averaging with a width of typically three time windows in between minimization steps. The final drift estimate however, is not smoothed. Finally, the drift estimate is interpolated to individual frames of the raw SMLM dataset.

##### Temporal Segmentation methods

Three distinct segmentation methods were implemented to segment SMLM datasets into time windows:

1. **Frames per segment**: Dataset is segmented into equally sized time windows based on the specified number of frames per segment.
2. **Localizations per segment (default)**: Dataset is segmented to contain an equal number of localizations per segment, ensuring uniform statistical significance across segments.

3. **Number of segments**: The dataset is divided into a specified total number of segments, automatically adjusting the number of localizations per segment.

To achieve homogeneous distribution of localizations, content-aware segmentation was used were applicable. Specifically, if the last segment contained fewer localizations than the segment average, it was merged with the preceding segment. Localizations acquired in the same frame were always assigned to the same time window. Whenever down sampling was necessary, it was performed randomly yet optimized to maintain homogeneous information content per segment. For datasets acquired with one-directional axial scanning, segmentation edges were explicitly aligned with z-plane boundaries, wrapping segments around accordingly.

#### Technical details

Pre-processing, segmentation, and localization pair identification were implemented using NumPy arrays.[1] To find the localization pairs, the KD-tree of the localization coordinates is created using the SciPy package,[2] using the 'query_pairs' function. The L-BFGS-B implementation was also provided by SciPy.

Computationally intensive cost-function and gradient (via derivative computations) evaluations are GPU-accelerated using CUDA via Numba's cuda.jit function to compile a custom code.[3] Specific software versions: Python 3.8, SciPy 1.7.3, Numba 0.55, CUDA toolkit 11.2, GPU driver version 460.

All benchmarking experiments were conducted on a workstation equipped with an Intel Xeon W-2223 CPU, 32 GB RAM, and an NVIDIA Quadro P2200 GPU.

### Simulations

#### Artificial drift generation

Artificial drift was generated to incorporate the key features often found in real drift comprising relaxation, random and oscillatory components,[4] here simulated using a combination of slow and fast oscillations as well as two exponential decay components. The explicit drift function used was:

$$\vec{D}(t) = \vec{A}_{slow} \sin\left(\frac{2\pi t}{\tau_{slow}} + \vec{\phi}_{slow}\right) + \vec{A}_{fast} \sin\left(\frac{2\pi t}{\tau_{fast}} + \vec{\phi}_{fast}\right) + \vec{A}_{d1} e^{-t/\tau_{d1}} + \vec{A}_{d2} e^{-t/\tau_{d2}}$$

The timescales $\tau_{fast}, \tau_{slow}$ of the oscillatory components used for the simulations were 200 and 6000 frames respectively. Furthermore, the amplitudes $\vec{A}$ were chosen to feature a small continuous change over time extracted from a smoothed random walk. The resulting drift curve can be seen in Supplementary Fig. 6.

#### Dataset resampling

To generate a test dataset, that by definition does not contain any drift, but still is as realistic as possible, a real SMLM datasets was resampled, treating every (grouped and filtered by usual means) localization in the existing dataset as a ground truth label site and creating a new dataset based upon this. Resampling procedures for these SMLM datasets included randomly selecting a set of - on average – $m$ label sites per newly generated frame over a fixed total number of frames, sampling their coordinates and, optionally, applying a normally distributed positional offset to simulate finite localization precision.

Tubulin Dataset. A 2D STORM dataset from a measurement on U2OS cells targeting Alpha-tubulin immuno-labelled with Alexa 647 was downloaded from Shareloc.xyz (10.5281/zenodo.7234161), resampled to a dataset containing 60k frames with 20 localizations per frame and the artificial drift applied as described previously. A rendering of the drift corrected dataset can be seen in Supplementary Fig. 10.

NPC Dataset. A 3D 4Pi-STORM dataset from a measurement on U2OS cells targeting the nuclear pore complex (Acquisition Details described in Bates et al.[4]) was resampled using 1.25 / 2 / 5 localizations per frame to three datasets of 60k frames resembling sparse, medium and rich conditions. A rendering of the drift corrected dataset can be seen in Fig. 4a.

#### Cost-function landscape investigations on resampled tubulin dataset

To gain insight into the optimization landscape underlying the COMET algorithm, we systematically investigated the cost-function as a function of two key control parameters: the Gaussian scale parameter $\sigma$ and the temporal segmentation granularity. Understanding how these parameters shape the landscape is essential, because the convergence behaviour of the optimizer and ultimately the accuracy of the drift estimate both depend on the smoothness and convexity of cost-function landscape in the vicinity of the global minimum. We first examined the dependence of $f_C$ on $\sigma$. The resampled tubulin dataset was drifted using the artificial drift trajectory, down-sampled twenty-fivefold after segmentation into single-frame time windows and paired within a 500 nm radius. The drift amplitude was then scaled continuously between −2.0 and +2.0 (in steps of 1%), and the cost function was evaluated at each point for $\sigma$ of 1, 2, 4, 16, 64, and 256 nm. The resulting one-dimensional slices through the high-dimensional cost-function landscape (Supplementary Fig. 7a) confirm that larger $\sigma$ values produce a smoother landscape with a single, broad global minimum, facilitating initial convergence of the optimizer. As $\sigma$ decreased, the landscape develops sharper features that encode finer spatial information, enabling progressive refinement of the drift estimate, the principle underlying the iterative $\sigma$-reduction protocol used by COMET.

To quantify the residual error of an ideal estimator operating on this landscape, a second series of calculations was performed at finer amplitude resolution (steps of 0.2%, covering amplitudes from −0.2 to +0.2) for $\sigma$ values of 1, 2, 4, 8, 16, 32, and 64 nm. For each $\sigma$, the minimum of $f_C$ was identified, the corresponding drift estimate was applied to the dataset, and the resulting localization accuracy was quantified as the RMS standard deviation relative to the ground-truth positions (Supplementary Fig. 7b).

We next assessed the influence of temporal segmentation on the cost-function landscape. $\sigma$ was fixed at 1 nm while the number of frames per time segment was varied over the values 1, 4, 16, 64, 256, and 1024. For each condition, the drift estimate was sub-sampled by averaging the ground-truth drift within each segment (Supplementary Fig. 7c curves offset by 50 nm for clarity). The corresponding cost-function slices (Supplementary Fig. 7d) reveal that coarser segmentation smooths the landscape at the expense of temporal detail, whereas finer segmentation preserves fast drift transients but increases the dimensionality of the optimization. A complementary fine-resolution scan (0.2% steps, amplitudes −0.2 to +0.2) for 1, 2, 4, 8, 16, 32, 64, 128, and 256 frames per window confirmed this trade-off and provided the localization-accuracy curves shown in Supplementary Fig. 7e.

#### Algorithm robustness and sensitivity analysis of COMET

A practical drift-correction algorithm must not only be accurate under ideal conditions but also robust to the choice of user-specified parameters. In the COMET framework, the two primary parameters exposed to the user are the maximum expected drift and the initial Gaussian scale parameter $\sigma$. To establish practical guidelines and verify that COMET performs reliably without extensive parameter tuning, we conducted a systematic sensitivity analysis on the resampled tubulin dataset. The aim of this analysis was twofold: first, to delineate the range of parameter choices over which COMET returns accurate drift estimates, and second, to confirm the heuristic that setting the initial $\sigma$ to approximately one-third of the maximum expected drift yields robust convergence across diverse conditions.

In the first set of experiments, the influence of the initial Gaussian scale parameter $\sigma$ was evaluated while keeping the maximum expected drift parameter fixed to the maximum of the simulated drift applied to the resampled dataset. The initial Gaussian scale parameter was set to 200, 150, 125, 100, 75, 60, 50, 40, 30 nm, with a target Gaussian scale parameter of 30 nm. For each condition tested, the accuracy of the drift estimate was quantified as the root mean square standard deviation (RMSD) from the to ground truth drift (Supplementary Fig. 8a). In the second set of experiments, the influence of the maximum drift parameter ($D_{max}$) was investigated while fixing the initial Gaussian scale parameter to one-third of the maximum drift, as suggested by the first set of experiments. The maximum drift parameter was explicitly set to 600, 500, 450, 400, 350, 300, 250, 200, 150, 100 nm and similarly as before using a target Gaussian scale parameter of 30 nm. As above, the estimated drift was compared to the ground truth, and the resulting errors were calculated (Supplementary Fig. 8b).

Taken together, these analyses demonstrate that COMET exhibits stable performance across a broad range of parameter choices. The recommended heuristic of Gaussian scale parameter to be 1/3 of the maximum expected drift with a target scale on the order of 10 nm proved sufficient to recover the drift to single-nanometer accuracy in all tested scenarios, confirming that careful but not exhaustive parameter selection is sufficient for reliable operation.

#### Optimal Segmentation for COMET on the resampled tubulin dataset

Beyond the Gaussian scale and maximum drift parameters, the temporal segmentation of the localization dataset, i.e., the number of localizations assigned to each time window, is the primary lever controlling the trade-off between temporal resolution and statistical reliability of the drift estimate. Too few localizations per segment yield an under-determined optimization problem, while too many average out fast drift transients. To identify the optimal operating point for a representative dataset, we performed a systematic segmentation sweep on the resampled tubulin dataset with the aim of quantifying this trade-off and providing a practical guideline for users. Using a maximum expected drift of 300 nm and the initial Gaussian scale parameter of 100 nm the dataset was segmenting using 200, 300, 400, 500, 600, 800, 1000, 1500 and 1750 localizations per segment. The boxcar smoothing was set to 3 timepoints in between the optimization steps. We characterized the performance by calculating the standard deviation of the difference in interpolated drift estimate and ground truth drift curve in X, Y and Z respectively which led to the datapoints displayed in Fig. 3e. The optimal solution we found was using 400 localizations per segment with an average error below 1 nm, with the resulting drift curve and residual difference to the ground truth per frame shown in Fig. 3c. These results establish that, for filamentous structures typical of cytoskeletal SMLM imaging, a segmentation of approximately 400 localizations per time window achieves sub-nanometer drift estimation accuracy. As a general guideline, a few hundred localizations per segment provides a robust starting point for most SMLM datasets, with the optimal value depending on the spatial complexity of the underlying sample structure.

#### Benchmarking with state-of-the-art software packages

Benchmarking comparisons explicitly included AIM, Minimum Entropy (ME), RCC (via SMAP and own implementation, using projection renderings), and DCC (also own implementation, using projection renderings). To compare the different drift correction methods a high-quality 3D 4Pi-STORM dataset was resampled using 1.25 / 2 / 5 localizations per frame (sparse, medium and rich) and the artificial drift per frame was applied as described previously. To ensure a fair comparison the segmentation into time windows and the frame-wise interpolation was done using self-implemented Python code. Using 480, 240, 120, 60, 48, 24, 12, 6, 2 frames per segment the dataset of each condition was segmented, equivalent to ca.: 2400 - 2.5 localizations on average per time window. Starting multiple rounds from the coarsest segmentation for each condition and method the dataset was drift corrected, interpolated using a cubic spline and saved until the drift estimation failed (diverging drift estimate or error). For each of the sampling conditions the segmentation condition that yielded the smallest deviation from the ground truth drift, quantified as the standard deviation of the difference between interpolated drift estimate and ground truth, was selected for comparison. For each sampling condition, the segmentation yielding the smallest deviation from the ground-truth drift—quantified as the standard deviation of the difference between the interpolated drift estimate and the ground truth—was selected for comparison. Performance was evaluated using two metrics: error per localization and error per label site. For the error per localization, the offset between each corrected localization coordinate and its ground-truth coordinate was computed independently in the X, Y and Z directions. Figure 4b shows the two-dimensional histogram of X–Y offsets for the respective methods.

For the error per label site, the absolute distance between each localization coordinate and its corresponding ground-truth label site was computed and averaged per label site. The resulting distributions are shown as one-dimensional histograms in Supplementary Fig. 9a–c for the sparse, medium and dense sampling conditions, respectively.

Parameters for each method are listed below; see the respective documentation for detailed descriptions:

- **COMET**: Gaussian scale parameter (initial: 100 nm, target: 10 nm), maximum drift (500 nm)
- **AIM**: IntersectD = 0.2, roiR = 3
- **ME**: coarseFramesPerBin=100, coarseSigma=[0.2,0.2,0.2]
- **RCC (SMAP)**: window pix=7, maxdrift nm=1000, pixrec nm=10
- **RCC (Own)**: pixel size 40 nm, Gaussian rendering width 80 nm, window size of fitted region: 15 pixel
- **DCC**: pixel size 40 nm, Gaussian rendering width 80 nm, window size of fitted region: 15 pixel

Complete tables of residual drift and computational times for each dataset condition (rich, medium, sparse) are included (see Supplementary Tables 1-3).

#### Experimental Data

To demonstrate the versatility of COMET across SMLM modalities, we applied the algorithm to experimental datasets acquired using 4Pi-STORM, OligoSTORM, MINFLUX, and conventional 2D STORM under controlled mechanical disturbance. These applications are described in the following subsections.

##### 4Pi-STORM imaging

Sample preparation and imaging for 4Pi-STORM datasets was carried out as described previously.[4] Multicolor datasets were measured using Alexa 647 and Cy5.5 as the fluorescent labels.

##### Experimental 4Pi-STORM dataset processing

As a proof-of-principle COMET was applied to a 3D 4Pi-STORM measurement and compared with results obtained by using a self-implemented 3D-RCC to estimate the drift. To obtain a baseline for the correction of this dataset the drift was estimated using RCC in a series of experiments in each step increasing the number of time windows, increasing the temporal resolution of the drift estimate until the algorithm breaks down. Starting from 4000 localizations per time window (corresponding 68 segments) decreased by a factor of two each step, down to 1000 localizations per window (corresponding 275 time windows), at which point a large discrepancy between to the previous drift estimates appeared. Using COMET with 300 nm max. drift, 30 nm initial and target Gaussian scale parameter the dataset was corrected using the same approach starting with 68 time windows the dataset was corrected with segmentations all the way to 50 localizations per time window. To assess the credibility of the obtained drift estimates with higher temporal resolution than the RCC equivalent a self-consistency check was performed, in which the dataset was split into two independent halves, one made up from localizations stemming from an odd and the other one from even frame, assuring independency by grouping localizations stemming from one on-event lasting multiple frames prior the split. COMET was used to find a drift estimate in each half to retrieve the discrepancy of the measure of error in the drift estimate.

To further investigate the influence of segmentation on the drift estimate for different methods another 4Pi-STORM measurement targeting the nuclear pore complex was investigated. For measuring the influence of the segmentation on COMET, the acquired dataset was split into time segments with five conditions using on average 1000, 500, 250, 100 and 50 localizations per segment and independently analysed using COMET. As a comparison, the same dataset was also corrected with projection RCC, we found 250, 500 and 1000 localizations per time segment applicable. The calculation times for three conditions using both COMET and RCC are shown in Supplementary Fig. 11a.

Furthermore, after applying the drift estimates of RCC and COMET under the 250 localizations per segment condition to the datasets, a 2D projection of the dataset was rendered, a structure showing a single nuclear pore complex unit with a high apparent labelling efficiency was cropped and the localizations were rendered in 3D using ChimeraX.[5] The 2D and 3D renderings are shown in Supplementary Fig. 11b and 11c respectively for both COMET and RCC (blue, yellow). Although the segmentation condition was identical the reconstructed structures look very different, with the result based on the COMET correction resembling more the expected structure of the nuclear pore complex, specifically the 8-fold symmetry of the Nup96 target.

Comparing the frame-wise interpolated drift estimates and their standard deviation calculated from the three conditions each, as visualized in the plot in Supplementary Fig. 11d, we see a similar trend for RCC and COMET, however, the deviation in the individual RCC estimates is partly more than an order of magnitude higher than the deviation of the COMET estimates, hinting that the RCC might have already been prone to noise in the cross-correlation function and therefore yielding an unstable drift estimate.

On smaller timescales, COMET finds subtle drifts with amplitudes of 10 nm, especially in the drift along the Z-axis, which RCC seems to miss, which could explain why the rendered image of the RCC does not show the NPC-subunits as clearly, ultimately limiting the effective resolution of the final reconstructed super-resolved image to above that length scale.

#### 2D STORM acquisition under controlled mechanical disturbance

We acquired 5,000 frames of 2D STORM data on a commercial Abbelight system (SAFe MN360) with an exposure time of 10 ms per frame during a live demonstration, with the active focus-stabilization system engaged. During the acquisition, we deliberately perturbed the microscope by applying two sequences of interleaved “strong” and “weak” pushes to the floating optical table supporting the instrument, in order to induce abrupt, large-amplitude drift. After each disturbance, the focus-stabilization system restored the correct focal plane within a few seconds, ensuring that the 2D imaging continued to probe the same underlying structure throughout the experiment.

#### Experimental processing of disturbed 2D STORM dataset

The acquired frames were analysed by the commercial Abbelight software (Abbelight NEO analysis). The dataset was segmented into approx. 1500 time windows using downsampling to reduce the data to approximately 250 localizations per time window. Using 1.8 µm as an upper bond for the maximum drift, ca. 1.26 billion pairs of localizations were passed into the COMET algorithm to estimate the drift. To refine the drift estimate even further, a refined COMET run was performed on the dataset corrected with the drift estimate from the first run using approx. 2440 segments using all localizations present in the dataset (on average 180 per time segment) and a maximum drift setting of 200 nm, this time yielding ca. 34 million pairs. The combination of the drift estimates of the first and second run yield the drift shown in Supplementary Fig. 13a. Fig. 13b and 13c show close-ups of regions in the drift estimate right after a disturbance was introduced, showing a periodic damped oscillation both in X and Y.

These regions were individually cut out and the period was measured to create the histogram of oscillation frequencies observed with a mean 3.1 Hz (Supplementary Fig. 13d).

#### Sequential OligoSTORM imaging

Genomic imaging with OligoSTORM[6] was performed on a commercial microscope, the Vutara VXL (Bruker Corp., USA), as described previously[6]. Utilizing PaintSHOP (paintshop.io), a set of probes targeting a 500 kb region on chromosome 2 (chr2:112.67–113.17 Mb, hg38) was designed with an Oligopaint probe (OP) density of 5.4 OP per kilobase (kb). Oligopaint barcode appending was performed using a custom script. The readout design for iterative fluorophore removal (toeing) and rehybridization, permitting repeated imaging of the same genomic region with a different set of fluorophores. Library and readout probe details and ordering files can be shared upon request. Each readout probe set, conjugated with two Alexa Fluor 647 fluorophores, was introduced via fluidics and defined as a hybridization time point. For each experiment, 3 hybridization time points were acquired. At each time point, a 4-µm axial volume was imaged in 100-nm steps (41 z-positions), with 1,000 frames recorded per z-plane. The volume was scanned 5 times, yielding 205,000 frames per hybridization time point. During the experiment, as switching buffer, we introduce a freshly prepared 2-Mercaptoethanol (BME) buffer in a 50mM TRIS in 2x SSC buffer with an oxygen scavenging system (final concentrations): 10% D-Glucose (Sigma-Aldrich, #G8270), 0.625 mM Sodium Chloride, 150 ug/mL of Catalase (Sigma-Aldrich, C40, ≥10,000 units/mg protein), 1.5 mg/mL Glucose Oxidase (Sigma-Aldrich #G2133-250KU), adjusted to pH 8.0. The fluorophores were excited with 9-10 kW/cm^2 laser intensity with a wavelength of 640 nm in wide-field. Additionally, we utilized increasing 405 nm activation from 0 W/cm2 in cycle 1 to 10 W/cm2 in cycle 5.

<u>Cell preparation:</u> Human retinal pigment epithelial-1 (RPE-1) were grown at 37°C + 5% CO2 in serum-supplemented (10% v/v) Dulbecco's Modified Eagle Medium (DMEM) (serum Gibco 10437; media Gibco 10564). The cells were also supplemented with 1% (v/v) MEM Non-Essential Amino Acids Solution (Gibco 11140050). Penicillin and streptomycin (Gibco 15070) were also added to the cell culture media to final concentrations of 50 units/ml and 50 µg/ml, respectively.

<u>Fluorescence In-Situ Hybridization Protocol:</u> FISH (fluorescence in-situ hybridization) was performed as described in Nir et al.[6] In brief, ibidi samples were incubated in 0.1N HCl for 5 minutes at RT, followed by two washes with 2x SSCT (2x Saline Sodium Citrate buffer with 0.1% Tween20). RNAse treatment (50 µL 2x SSCT with 2 µL RNAse A) was applied for 60 minutes at 37°C. The samples were treated with 50% formamide (FA) in 2x SSCT for 10 minutes at RT, then 20 minutes at 60°C. Denaturation of the DNA was performed for 3 minutes at 80°C in a hybridization buffer containing the 100 pmol of the Oligoprobe library. Afterward, the samples were hybridized for > 48 hours at 42°C in a humidified chamber. Samples were washed twice with prewarmed 2x SSCT at 60°C for 10 minutes, followed by two additional washes at RT for 2 minutes. Fiducials were introduced by adding 40 µL of sonicated Tetraspeck (1:400 dilution in PBS of Tetraspeck) and gold nano-urchins (1:25 in PBS of gold nano-urchins,

GNUs; d = 90 nm, 630 nm abs max - Cytodiagnostics, Ontario, Canada) solution to the samples and centrifuging at 500g for 3 minutes. After rinsing with 2x SSCT and a final wash with 50% formamide in 2x SSCT, the samples underwent a 10-minute hybridization with the read-out probes in 40% formamide in 2x SSCT using a fluidics system. This was followed by a 10-minute wash in 50% formamide in 2x SSCT, then another 10-minute wash in 2x SSCT. Finally, the imaging buffer was introduced, and imaging was started. For time-point > 2, a toeing step is performed before the hybridization (10 min incubation of secondary probes with the appropriate toeing sequence in 40% FA in 2x SSCT).

#### Experimental OligoSTORM dataset localization and processing

Localization is performed with the vendor software SRX (v7.0.9 Bruker Corp USA) determining axial positions of the signal from the relative intensity and spatial distribution of the reference PSF measured on the same coverslip during calibration. User defined fitting parameters included a 16 px cutout window and a background threshold of BG50, which defines the signal-to-noise cutoff for event detection via bandpass filtering. Post-localization filtering removed events with axial precision below 20 nm in xy and 40 nm in z, with precision estimated using the Cramér–Rao lower bound. The fiducial-based drift correction was performed by selecting the area around fiducial markers (GNUs) manually in SRX. Fiducial localizations were filtered with more stringent precision thresholds (≤8 nm in *xy*, ≤16 nm in *z*), and a subset of 11 beads in this example was used to register via cross-correlation (Bruker proprietary algorithm) with a temporal segmentation of 100 frames per interval. The resulting drift correction trajectory (Fig. 5b,c) was applied to the full dataset. Prior to COMET drift estimation, the raw OligoSTORM localizations were pre-filtered: (i) localizations originating from fiducial beads were removed, and (ii) localizations in the extreme focal planes were discarded, as they contributed negligibly to the dataset. After filtering, each dataset was subsampled by hybridization time point, so that drift estimation was performed independently for each time point. Each hybridization time dataset was segmented into 250 time windows, yielding approximately 600 localizations per COMET-defined time segment, and the COMET drift estimate was then calculated independently. To reconstruct the global drift trajectory across multiple hybridization time points, the time point–specific drift estimates were aligned. Alignment was performed by estimating the relative translational shift between rendered localization histograms of individual time points using cross-correlation. To compare the COMET result with the fiducial-based drift correction, the same initial filtering steps were applied. The registered and filtered localization tables were then exported to CIMA (https://gitlab.iit.it/ina/CIMA) for signal detection, assessment of chromatin segments, and comparison.

#### MINFLUX Imaging

MINFLUX nanoscopy achieves localization precisions in the single-nanometer regime, making it one of the most demanding SMLM modalities with respect to drift correction. Even residual drift of a few nanometers can substantially degrade MINFLUX image quality. Commercially available MINFLUX instruments typically include hardware-based stabilization systems using fiducial tracking; however, residual drift may persist, and post-acquisition algorithmic correction can further improve the result. We therefore applied COMET to a MINFLUX DNA-PAINT dataset to evaluate whether the algorithm can operate effectively in this high-precision regime and provide a meaningful improvement over hardware stabilization alone. A Gattaquant 3x80 nm nanoruler sample was imaged on commercial MINFLUX (Abberior Instruments) using a standard 2D imaging sequence. DNA-PAINT MINFLUX using Cy3b imager strands in Massive Photonics imaging buffer and 560 nm excitation. 150 nm gold beads were added to the sample as fiducial markers, enabling continuously stabilization of the sample of approx. 1 nm in X, Y and Z by the built-in stabilization system.

##### Experimental MINFLUX dataset processing

The exported dataset was analysed, grouping localizations from the same trace (consecutive localizations during one binding event) and determining their mean position. The dataset was then segmented into time windows consisting of 25 traces per time window. The maximum expected drift was estimated from the raw localization histogram to be less than 150 nm. Raw and drift corrected rendered localizations as well as the COMET drift estimate are shown in Supplementary Fig. 13a,b,c and d respectively. The drift-corrected MINFLUX data show a visibly crisper point clouds, confirming that COMET can resolve and correct residual drift at length scales relevant to MINFLUX imaging, even in the presence of active hardware stabilization.

#### Further Information and data availability

A paper repository containing all analysis scripts and means to reproduce the figures shown in this manuscript is published (https://github.com/gpufit/Comet). Files which are not included in the repository (due to storage limitations) are available upon reasonable request.

In the accompanying repository, clear instructions are provided for:

- Installing COMET: Standalone and integration into SMLM software (e.g. Picasso)
- Cloud-based execution (Google Colab notebook, dedicated COMET server)
- Input/output formats, workflows, examples
- Explicit code snippets demonstrating typical COMET usage scenarios.

Furthermore, the tested dataset for the benchmarking comparison, the applied simulated drift, the respective drift estimates as well as scripts used to generate them are provided in the accompanying repository.

### Software Implementation and Usage Notes

#### Choosing the Right Version of COMET

COMET is available in four different formats, each suited to different levels of expertise and use cases:

1. COMET Website
   The COMET web interface is designed for maximum accessibility and convenience. It allows users to quickly correct individual datasets in the standard ThunderSTORM CSV format, providing corrected results within minutes without hardware setup or software installation required. This is the ideal choice for users seeking a simple, no-code solution.
2. Google Colab Notebook
   The Colab version is best suited for users who don't have local GPU hardware but want more flexibility than the website offers. With basic knowledge of Python, users can follow the drift correction process step by step, examine the underlying code, and even adapt the notebook to handle non-standard file formats. Integration with Google Drive also enables batch processing of multiple datasets.
3. Picasso Integration
   For users already working with the Picasso software suite for SMLM data analysis, COMET can be installed as an available drift correction option. This version offers a seamless workflow and serves as a practical example of how COMET can be integrated into existing analysis pipelines.
4. COMET Source Code
   For advanced users with Python experience, the full COMET source code offers complete flexibility. It enables custom integration into bespoke analysis pipelines, supports arbitrary data formats and segmentation schemes, and includes all tools necessary to reproduce the results from the COMET publication. The source code repository includes example scripts and detailed documentation for adapting COMET to specific experimental scenarios.

If unsure where to start, the website is the quickest entry point, while the Colab notebook offers a balanced path between flexibility and ease of use. Advanced users can explore the source code for full integration into complex workflows.

#### Additional Notes and Recommendations

- **Data format compatibility**: All versions of COMET expect localization data with at least two-dimensional coordinates (X and Y) with a timestamp per coordinate (typically frames).
- **GPU requirements**: While GPU acceleration is essential especially for large datasets (typ. ≥2GB VRAM), the Colab and web versions provide cloud-based GPU support for users without local hardware.
- **Batch processing**: For correcting multiple datasets, we recommend using either the Colab version (via Google Drive integration) or the source code, which provides more control over scripting and automation.
- **Parameter Choice**: A few hundred localizations per time window is usually a robust choice for the segmentation, the drift estimate can easily be refined in consecutive runs using less localizations per time window until convergence is reached, or the drift curve changes significantly (break down of the method). The choice of the maximally expected drift is crucial, underestimation can lead to flawed results, while a strong overestimation significantly slows down the drift estimation process and can lead to unnecessary high hardware requirements. Usually, a visual inspection of a rendering of the raw localizations is enough to estimate the drift from the common spread of the visible point clouds. The initial Gaussian scale parameter should be set to 1/3 of the maximum drift and the target scale must not be smaller than the expected intrinsic localization precision.

### Supplementary Figures

#### Supplementary Figure 1

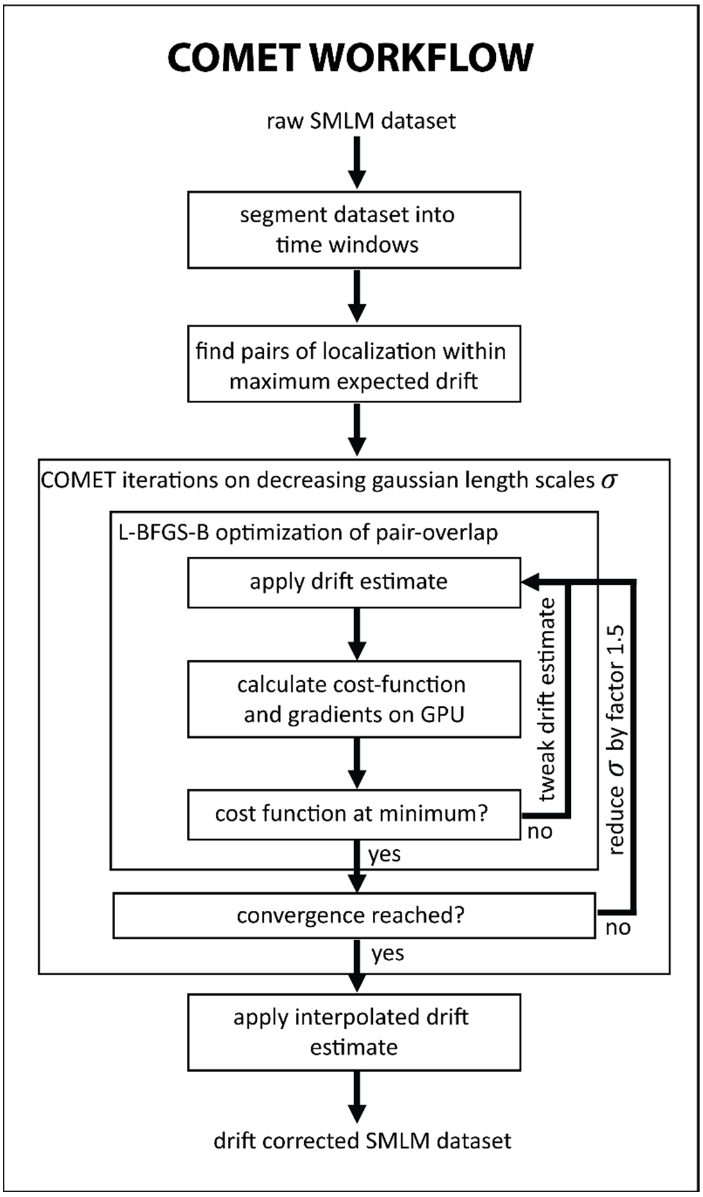

**Supplementary Figure 1: COMET algorithm workflow.** Flowchart illustrating the processing pipeline of the COMET drift correction algorithm. A raw SMLM dataset is first segmented into temporal windows and localization pairs within the maximum expected drift distance are identified. The core optimization loop iteratively minimizes the COMET cost function using the L-BFGS-B algorithm, with the Gaussian interaction scale sigma reduced by a factor after each successful convergence step. The procedure terminates when further reduction of sigma yields no improvement in the drift estimate. The final drift trajectory is interpolated to individual acquisition frames and applied to the localization coordinates.

Supplementary Figure 2

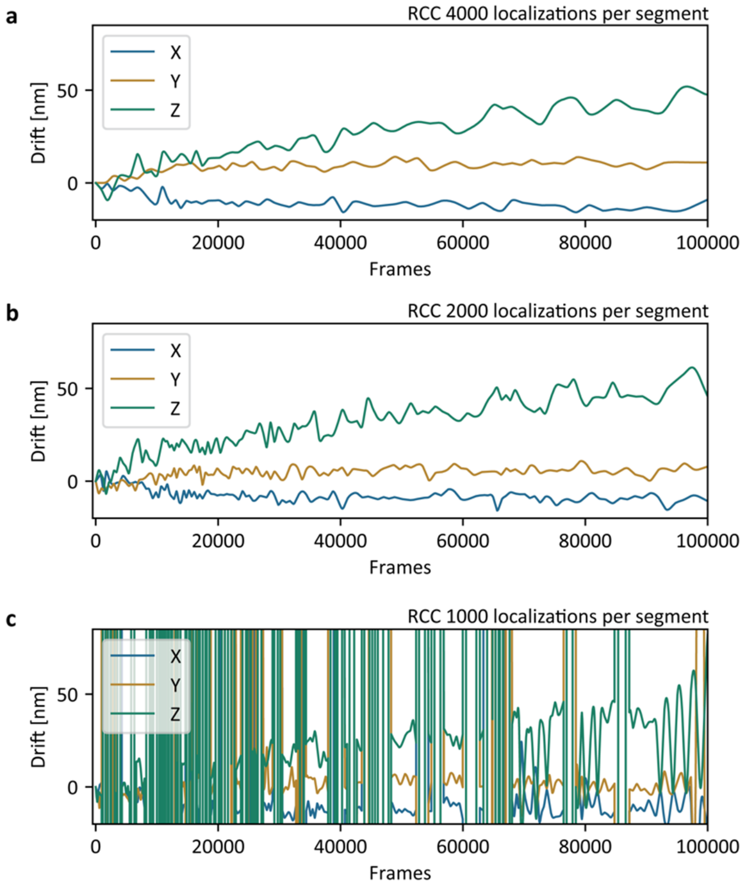

**Supplementary Figure 2: RCC drift estimates with increasing temporal resolution**. Three-dimensional drift estimates obtained by redundant cross-correlation (RCC) applied to an experimental 4Pi-STORM dataset of nuclear pore complexes, using (a) 4000, (b) 2000, and (c) 1000 localizations per time segment. Drift in X (blue), Y (orange), and Z (green) is plotted as a function of acquisition frame number. At 4000 localizations per segment the RCC solution is stable, but increasing the temporal resolution to 1000 localizations per segment leads to divergent drift estimates and spurious outliers, indicating the practical resolution limit of the method on this dataset.

#### Supplementary Figure 3

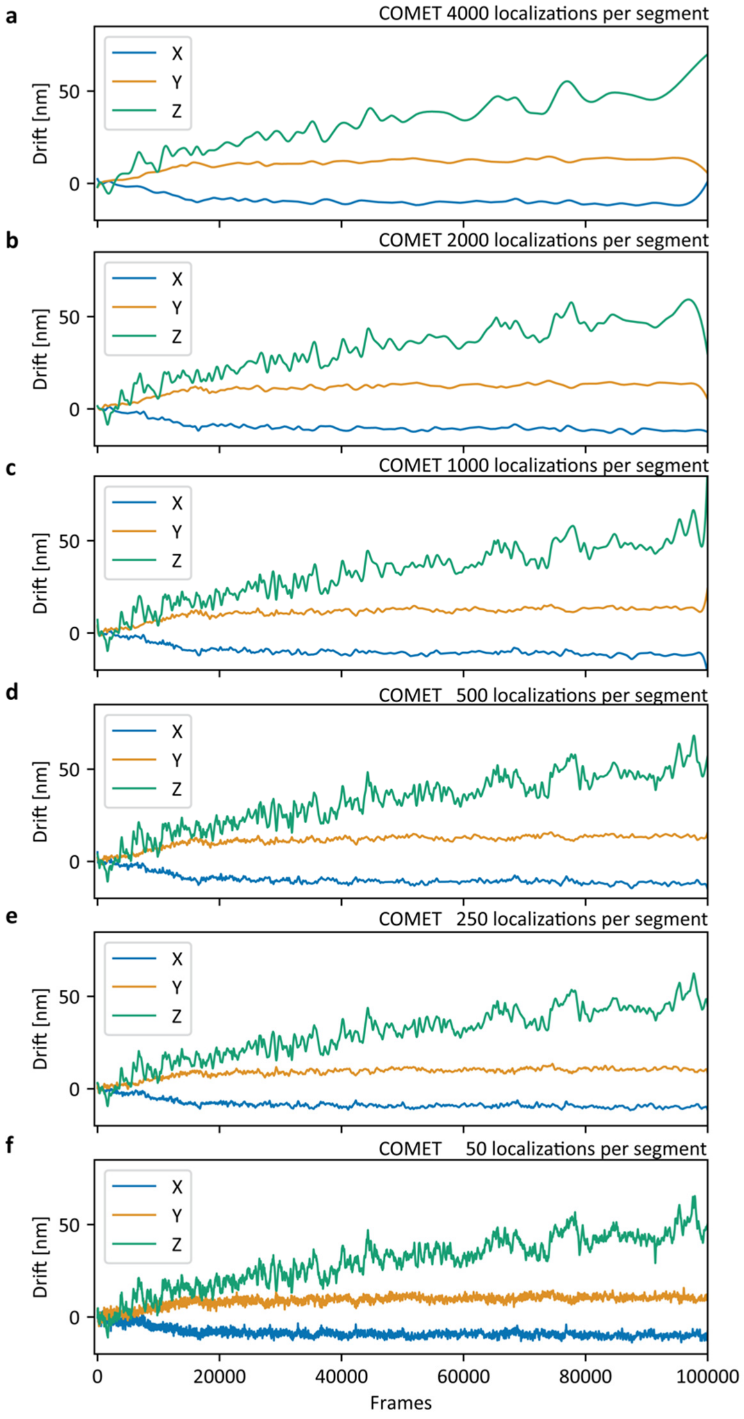

**Supplementary Figure 3: COMET drift estimates with increasing temporal resolution**. Drift trajectories estimated by COMET on the same experimental 4Pi-STORM NPC dataset as in Supplementary Figure 2, using (a) 4000, (b) 2000, (c) 1000, (d) 500, (e) 250, and (f) 50 localizations per time segment. X (blue), Y (orange), and Z (green) drift components are shown. In contrast to RCC (Supplementary Figure 2), COMET produces stable drift estimates across all tested segment sizes. Progressively finer temporal segmentation reveals fast drift transients that are masked at coarser resolutions, while the overall drift trajectory remains self-consistent.

Supplementary Figure 4

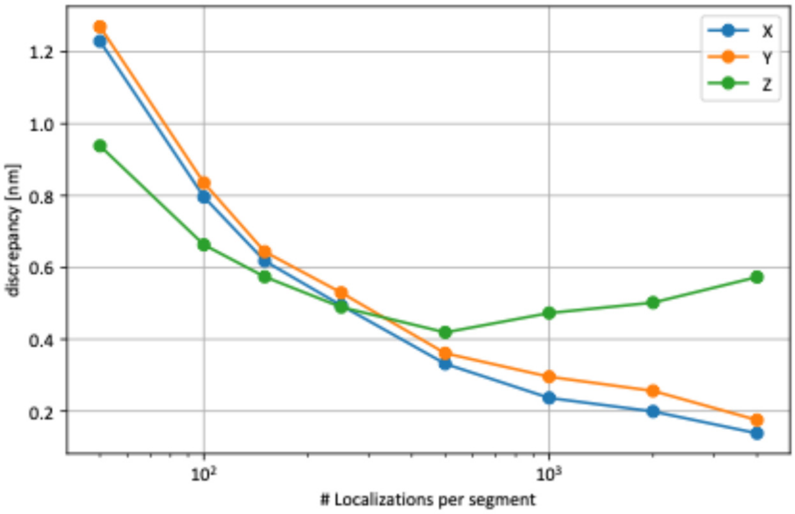

**Supplementary Figure 4: Self consistency of COMET drift estimates for different segmentation conditions.** The discrepancy of the COMET drift estimates when processing either only localizations from odd or even frames is calculated for X, Y and Z respectively for the tested segmentation conditions ranging from 50 to 4000 localizations per time window.

Supplementary Figure 5

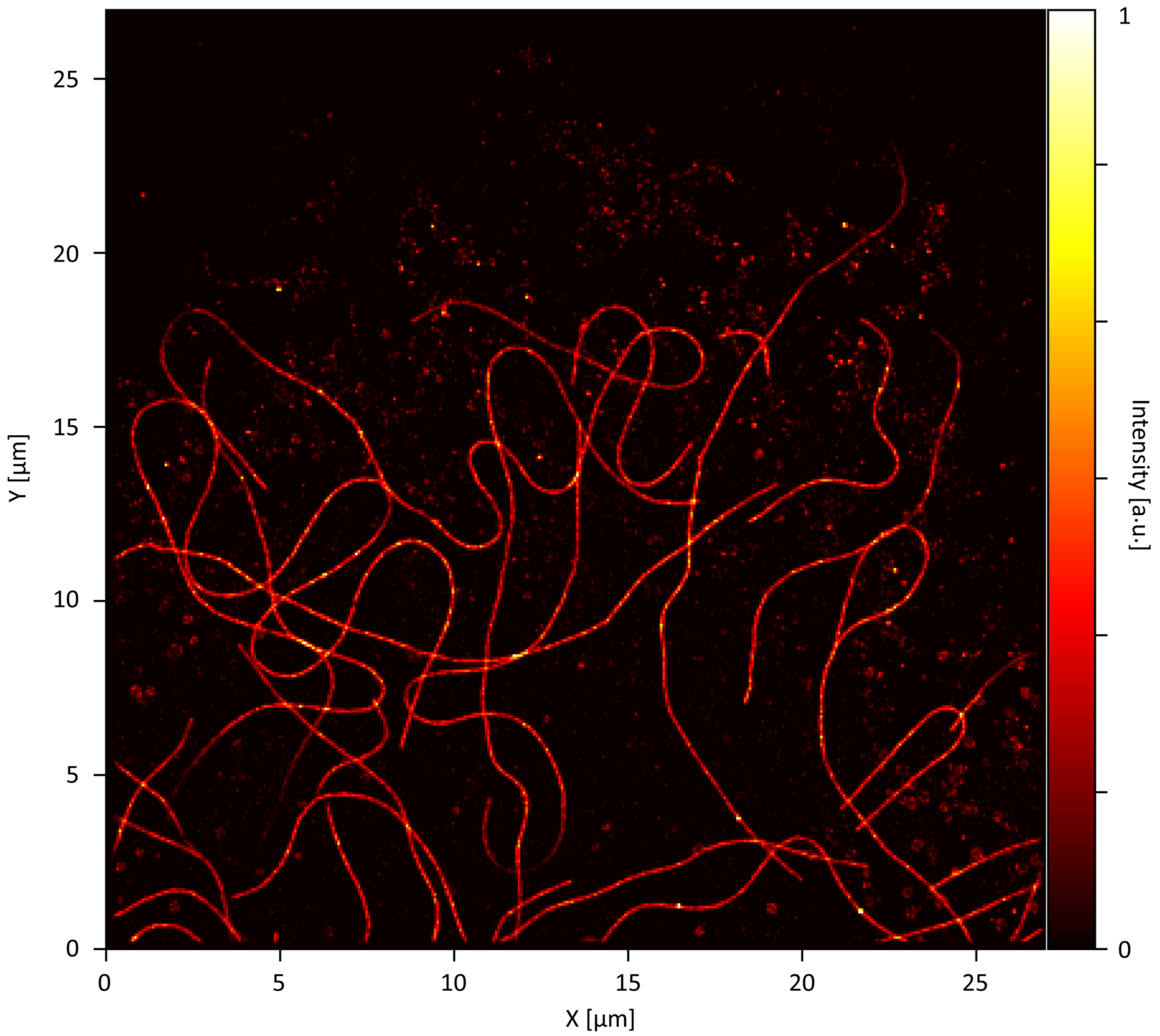

**Supplementary Figure 5: SMLM rendering of the resampled tubulin dataset after drift correction**. Super-resolution image of alpha-tubulin in U2OS cells, reconstructed from a 2D STORM dataset downloaded from ShareLoc (10.5281/zenodo.7234161) and resampled to 60,000 frames with 20 localizations per frame. The image shows the dataset after application of the COMET drift correction. Colormap indicates localization density. This resampled dataset served as the basis for the simulated drift experiments and benchmarking analyses presented in Figures 3 and 4 of the main text.

Supplementary Figure 6

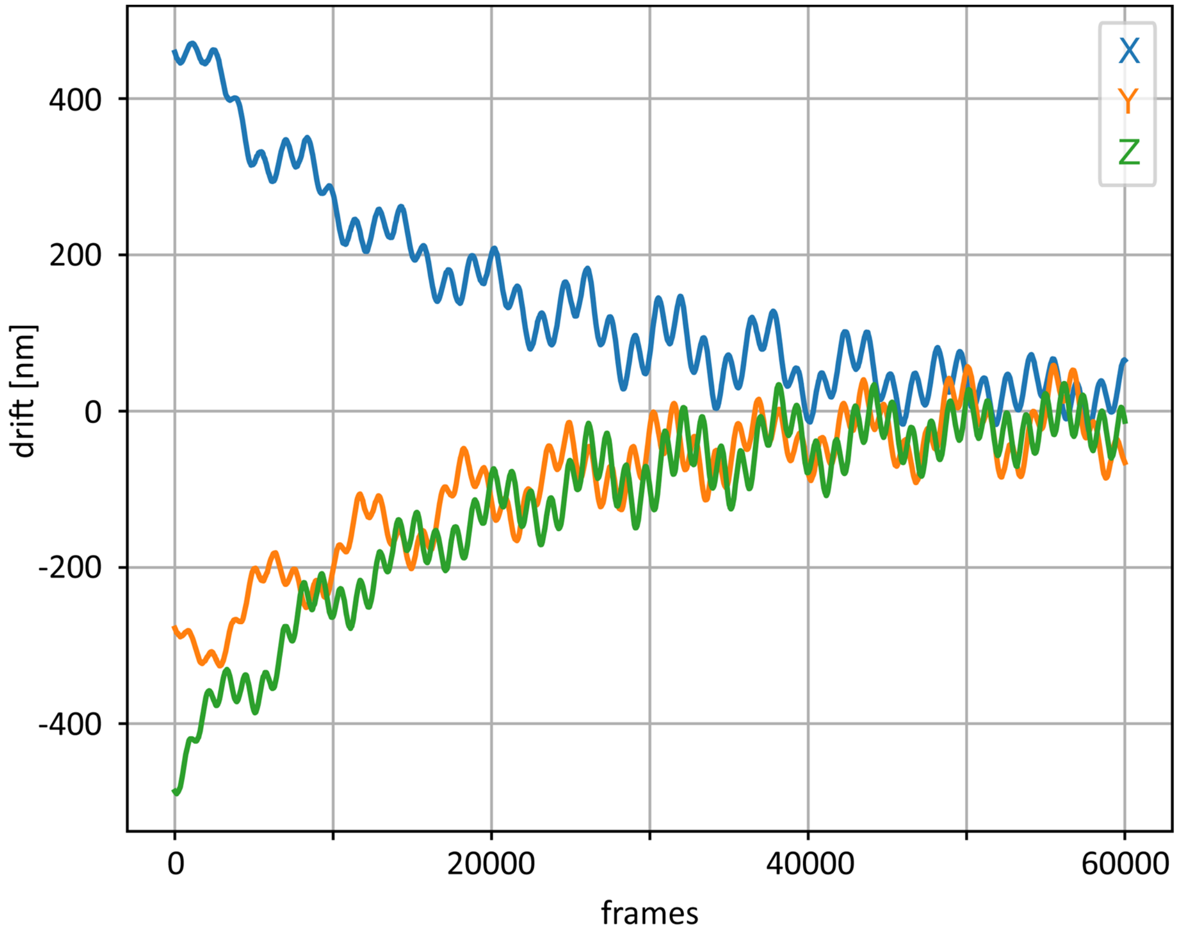

**Supplementary Figure 6 : Simulated three-dimensional drift trajectory.** Artificial drift applied to the resampled tubulin dataset, plotted as a function of acquisition frame for the X (blue), Y (orange), and Z (green) dimensions. The drift function comprises slow and fast sinusoidal oscillations superimposed on two exponential decay components, approximating experimentally observed drift dynamics that include both relaxation and periodic features. See Supplementary Methods for the explicit functional form and parameter values.

Supplementary Figure 7

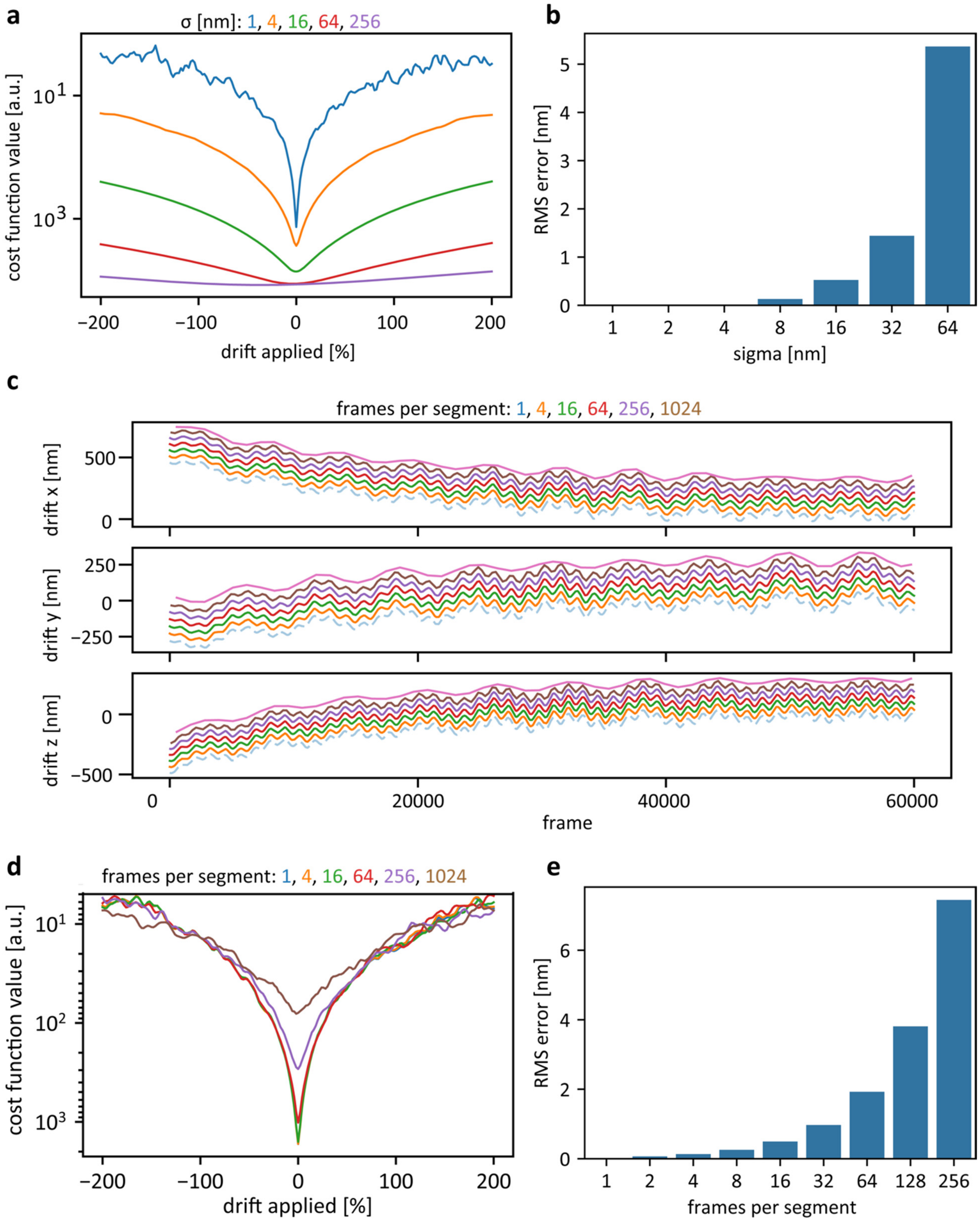

**Supplementary Figure 7: Cost-function landscape analysis of COMET.** (a) One-dimensional slices through the COMET cost function, computed by applying scaled versions of the simulated drift (amplitude factor from -2.0 to +2.0) to the resampled tubulin dataset, for Gaussian interaction scales sigma = 1, 4, 16, 64, and 256 nm. Larger sigma values produce smoother landscapes that facilitate convergence toward the global minimum; smaller sigma values sharpen the minimum, enabling higher-precision drift estimation. (b) Root-mean-square (RMS) error of an ideal drift estimator (defined by the cost-function minimum) as a function of sigma, quantified as the standard deviation of the distance between corrected and ground-truth localization positions. (c) Sub-sampled drift estimates for different numbers of frames per time window (1, 4, 16, 64, 256, 1024), offset by 50 nm each for visual clarity. The drift in X, Y, and Z is shown across 60,000 frames.(d) Cost-function landscapes for varying temporal segmentation (1, 4, 16, 64, 256, and 1024 frames per segment) at a fixed sigma = 1 nm. (e) RMS error of the ideal estimator as a function of frames per segment, showing increasing error with coarser temporal resolution. C

Supplementary Figure 8

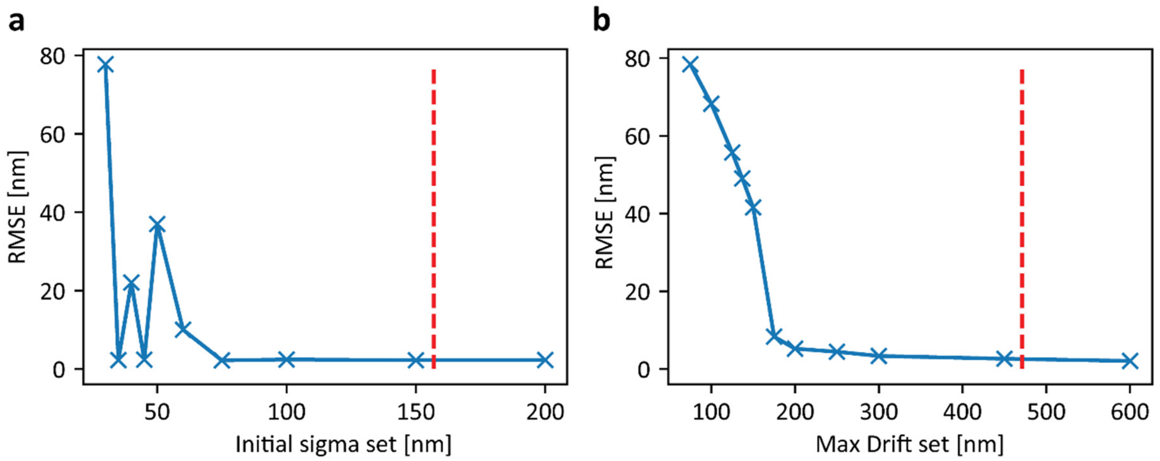

**Supplementary Figure 8: Parameter sensitivity analysis of COMET.** (a) Root-mean-square error (RMSE) of the COMET drift estimate on the resampled tubulin dataset as a function of the initial Gaussian interaction scale sigma, with the maximum expected drift fixed at the true maximum of the applied simulated drift (dashed red line indicates sigma = D_max / 3). Target sigma was set to 30 nm in all runs. The algorithm converges robustly for initial sigma values above approximately 60 nm. (b) RMSE as a function of the maximum expected drift parameter D_max, with the initial sigma set to D_max / 3. The dashed red line indicates the true maximum drift amplitude. Performance degrades sharply when D_max strongly underestimates the actual drift, but remains stable across a broad range. Together, these results confirm that COMET is robust to parameter choices within the recommended operating regime.

### Supplementary Table 1

| frames/bin | 480 | 240 | 120 | 60 | 48 | 24 | 12 | 6 |
|---|---|---|---|---|---|---|---|---|
| eq. # segment | 125 | 250 | 500 | 1000 | 1250 | 2500 | 5000 | 10000 |
| localizations/window | 2400 | 1200 | 600 | 300 | 240 | 120 | 60 | 30 |
| **COMET** | | | | | | | | |
| Residual drift X [nm] | 9,15 | 1,44 | 0,57 | 0,36 | 0,34 | 0,36 | 0,45 | 5,72 |
| Residual drift Y [nm] | 9,34 | 1,40 | 0,52 | 0,33 | 0,31 | 0,34 | 0,44 | 4,53 |
| Residual drift Z [nm] | 12,17 | 1,82 | 0,67 | 0,37 | 0,35 | 0,36 | 0,45 | 1,52 |
| calc time [s] | 48,15 | 20,22 | 18,48 | 20,09 | 19,51 | 28,07 | 36,05 | 41,10 |
| mean std [nm] | 10,22 | 1,55 | 0,59 | 0,35 | 0,33 | 0,35 | 0,45 | 3,92 |
| **Min. Entropy** | | | | | | | | |
| Residual drift X [nm] | 4,94 | 2,98 | 2,53 | 2,96 | 3,20 | 4,18 | 5,84 | 8,40 |
| Residual drift Y [nm] | 5,49 | 2,88 | 2,57 | 3,01 | 3,29 | 4,06 | 5,73 | 8,23 |
| Residual drift Z [nm] | 9,53 | 2,63 | 2,80 | 3,83 | 4,29 | 6,15 | 8,77 | 12,80 |
| calc time [s] | 66,47 | 64,23 | 68,68 | 86,19 | 90,01 | 76,71 | 96,67 | 89,40 |
| mean std [nm] | 6,65 | 2,83 | 2,64 | 3,27 | 3,59 | 4,80 | 6,78 | 9,81 |
| **AIM** | | | | | | | | |
| Residual drift X [nm] | 10,43 | 4,07 | 2,69 | 5,00 | | | | |
| Residual drift Y [nm] | 11,62 | 4,26 | 2,63 | 4,61 | | | | |
| Residual drift Z [nm] | 23,16 | 7,59 | 2,66 | 16088,77 | | | | |
| mean std [nm] | 15,07 | 5,31 | 2,66 | 5366,13 | | | | |
| calc time [s] | 0,75 | 1,21 | 1,87 | 3,56 | | | | |
| **RCC (SMAP)** | | | | | | | | |
| Residual drift X [nm] | 25,59 | 21,53 | 12,19 | 3,50 | | | | |
| Residual drift Y [nm] | 24,58 | 19,64 | 12,58 | 3,72 | | | | |
| Residual drift Z [nm] | 27,67 | 24,04 | 22,20 | 15,73 | | | | |
| mean std [nm] | 25,95 | 21,74 | 15,66 | 7,65 | | | | |
| calc time [s] | 480,00 | 188,00 | 753,00 | 3355,00 | | | | |
| **RCC (2D projections)** | | | | | | | | |
| Residual drift X [nm] | 4,90 | 2,90 | 4,10 | 5,90 | | | | |
| Residual drift Y [nm] | 5,80 | 2,50 | 3,60 | 5,40 | | | | |
| Residual drift Z [nm] | 8,50 | 3,20 | 4,00 | 5,70 | | | | |
| mean std [nm] | 6,40 | 2,87 | 3,90 | 5,67 | | | | |
| calc time [s] | 258,00 | 1033,00 | 4102,00 | 13508,00 | | | | |
| **DCC (2D projections)** | | | | | | | | |
| Residual drift X [nm] | 13,40 | 55,00 | 82,80 | | | | | |
| Residual drift Y [nm] | 10,80 | 13,90 | 110,20 | | | | | |
| Residual drift Z [nm] | 9,40 | 6,10 | 15,60 | | | | | |
| mean std [nm] | 11,20 | 25,00 | 69,53 | | | | | |
| calc time [s] | 12,10 | 25,00 | 54,10 | | | | | |

**Supplementary Table 1: Benchmark matrix rich condition.**

### Supplementary Table 2

| frames/bin | 480 | 240 | 120 | 60 | 48 | 24 | 12 | 6 |
|---|---|---|---|---|---|---|---|---|
| eq. # segment | 125 | 250 | 500 | 1000 | 1250 | 2500 | 5000 | 10000 |
| localizations/window | 960 | 480 | 240 | 120 | 96 | 48 | 24 | 12 |
| **COMET** | | | | | | | | |
| Residual drift X [nm] | 10,36 | 2,04 | 0,77 | 0,48 | 0,50 | 0,55 | 0,75 | 16,57 |
| Residual drift Y [nm] | 10,90 | 2,02 | 0,77 | 0,46 | 0,47 | 0,53 | 0,75 | 13,74 |
| Residual drift Z [nm] | 16,66 | 2,82 | 0,95 | 0,52 | 0,52 | 0,54 | 0,73 | 12,70 |
| calc time [s] | 9,29 | 4,97 | 4,63 | 4,50 | 5,94 | 6,82 | 10,23 | 34,30 |
| mean std [nm] | 12,64 | 2,29 | 0,83 | 0,49 | 0,50 | 0,54 | 0,74 | 14,34 |
| **Min. Entropy** | | | | | | | | |
| Residual drift X [nm] | 6,86 | 3,72 | 3,20 | 4,14 | 4,49 | 6,30 | 8,59 | 12,79 |
| Residual drift Y [nm] | 15,34 | 3,75 | 3,19 | 4,13 | 4,45 | 6,08 | 8,73 | 12,78 |
| Residual drift Z [nm] | 17,18 | 3,63 | 4,23 | 5,69 | 6,50 | 9,42 | 13,28 | 19,43 |
| calc time [s] | 14,41 | 13,36 | 13,95 | 14,89 | 15,73 | 16,39 | 17,68 | 21,10 |
| mean std [nm] | 13,13 | 3,70 | 3,54 | 4,65 | 5,15 | 7,26 | 10,20 | 15,00 |
| **AIM** | | | | | | | | |
| Residual drift X [nm] | 13,76 | 5,51 | 6,41 | | | | | |
| Residual drift Y [nm] | 17,74 | 5,75 | 6,94 | | | | | |
| Residual drift Z [nm] | 23,22 | 12,31 | 6946,35 | | | | | |
| mean std [nm] | 18,24 | 7,85 | 2319,90 | | | | | |
| calc time [s] | 0,35 | 0,56 | 1,02 | | | | | |
| **RCC (SMAP)** | | | | | | | | |
| Residual drift X [nm] | 26,89 | 21,69 | 9,80 | 4,00 | | | | |
| Residual drift Y [nm] | 27,90 | 20,15 | 12,57 | 4,10 | | | | |
| Residual drift Z [nm] | 30,81 | 25,16 | 23,02 | 12,43 | | | | |
| mean std [nm] | 18,24 | 22,34 | 15,13 | 6,84 | | | | |
| calc time [s] | 37,00 | 144,00 | 559,00 | 2369,00 | | | | |
| **RCC (2D projections)** | | | | | | | | |
| Residual drift X [nm] | 5,30 | 4,50 | 6,70 | 71,60 | | | | |
| Residual drift Y [nm] | 6,50 | 4,30 | 6,40 | 68,10 | | | | |
| Residual drift Z [nm] | 8,80 | 5,10 | 6,60 | 9,10 | | | | |
| mean std [nm] | 6,87 | 4,63 | 6,57 | 49,60 | | | | |
| calc time [s] | 242,00 | 1023,00 | 4341,00 | 18427,0 | | | | |
| **DCC (2D projections)** | | | | | | | | |
| Residual drift X [nm] | 34,30 | 57,40 | | | | | | |
| Residual drift Y [nm] | 47,70 | 35,20 | | | | | | |
| Residual drift Z [nm] | 17,50 | 9,50 | | | | | | |
| mean std [nm] | 33,17 | 34,03 | | | | | | |
| calc time [s] | 11,40 | 26,80 | | | | | | |

**Supplementary Table 2: Benchmark matrix medium condition.**

### Supplementary Table 3

| frames/bin | 480 | 240 | 120 | 60 | 48 | 24 | 12 | 6 |
|---|---|---|---|---|---|---|---|---|
| eq. # segment (ideal) | 125 | 250 | 500 | 1000 | 1250 | 2500 | 5000 | 10000 |
| localizations/window | 600 | 300 | 150 | 75 | 60 | 30 | 15 | 7,5 |
| **COMET** | | | | | | | | |
| Residual drift X [nm] | 11,56 | 5,00 | 1,06 | 0,67 | 0,64 | 0,75 | 7,05 | |
| Residual drift Y [nm] | 8,67 | 4,73 | 0,95 | 0,64 | 0,64 | 0,76 | 5,64 | |
| Residual drift Z [nm] | 24,03 | 6,18 | 1,19 | 0,81 | 0,70 | 0,76 | 5,89 | |
| calc time [s] | 5,24 | 5,07 | 2,49 | 3,00 | 3,11 | 4,78 | 7,99 | |
| mean std [nm] | 14,75 | 5,30 | 1,07 | 0,71 | 0,66 | 0,76 | 6,19 | |
| **Min. Entropy** | | | | | | | | |
| Residual drift X [nm] | 13,15 | 5,03 | 3,58 | 4,61 | 5,11 | 7,01 | 10,06 | 15,07 |
| Residual drift Y [nm] | 12,74 | 4,47 | 3,82 | 4,82 | 5,24 | 7,14 | 10,01 | 14,75 |
| Residual drift Z [nm] | 21,61 | 4,16 | 4,78 | 6,66 | 7,62 | 10,80 | 15,47 | 22,87 |
| calc time [s] | 9,47 | 7,88 | 8,19 | 8,97 | 8,85 | 9,27 | 9,83 | 11,03 |
| mean std [nm] | 15,83 | 4,55 | 4,06 | 5,36 | 5,99 | 8,32 | 11,85 | 17,56 |
| **AIM** | | | | | | | | |
| Residual drift X [nm] | 18,95 | 6,90 | 52,58 | | | | | |
| Residual drift Y [nm] | 18,21 | 7,05 | 57,27 | | | | | |
| Residual drift Z [nm] | 36,61 | 23,29 | 6138,12 | | | | | |
| mean std [nm] | 24,59 | 12,41 | 2082,66 | | | | | |
| calc time [s] | 0,24 | 0,38 | 0,75 | | | | | |
| **RCC (SMAP)** | | | | | | | | |
| Residual drift X [nm] | 29,69 | | | | | | | |
| Residual drift Y [nm] | 26,58 | | | | | | | |
| Residual drift Z [nm] | 33,61 | | | | | | | |
| mean std [nm] | 29,96 | | | | | | | |
| calc time [s] | 30,81 | | | | | | | |
| **RCC (2D projections)** | | | | | | | | |
| Residual drift X [nm] | 6,10 | 5,80 | 8,70 | | | | | |
| Residual drift Y [nm] | 7,60 | 6,00 | 8,80 | | | | | |
| Residual drift Z [nm] | 8,80 | 5,90 | 8,40 | | | | | |
| mean std [nm] | 7,50 | 5,90 | 8,63 | | | | | |
| calc time [s] | 241,00 | 1060,00 | 5425,00 | | | | | |
| **DCC (2D projections)** | | | | | | | | |
| Residual drift X [nm] | 26,20 | 143,00 | | | | | | |
| Residual drift Y [nm] | 30,80 | 86,10 | | | | | | |
| Residual drift Z [nm] | 23,00 | 21,00 | | | | | | |
| mean std [nm] | 26,67 | 83,37 | | | | | | |
| calc time [s] | 9,96 | 21,70 | | | | | | |

**Supplementary Table 3: Benchmark matrix sparse condition.**

### Supplementary Figure 9

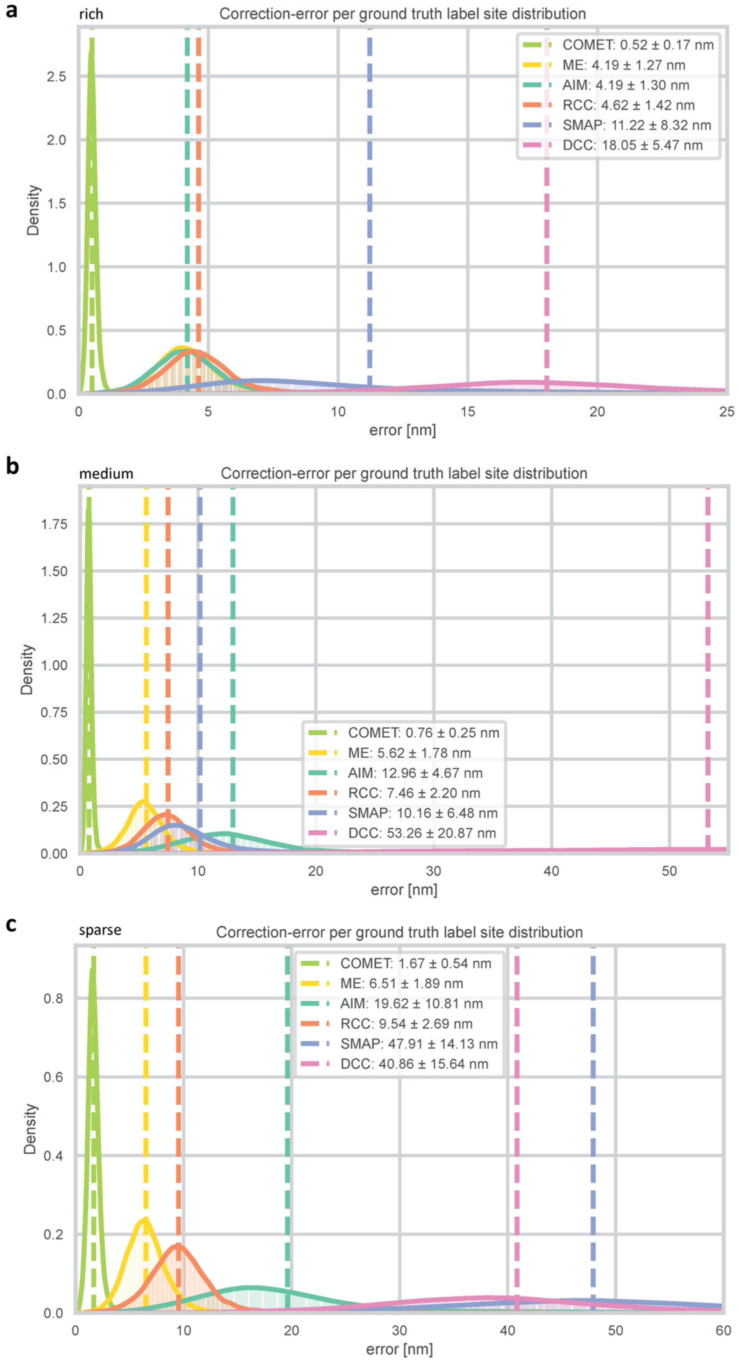

**Supplementary Figure 9: Benchmarking: correction error per ground-truth label site.** Histograms of the mean distance between drift-corrected localizations and their corresponding ground-truth label sites, computed for each drift correction method under (a) rich (5 loc./frame), (b) medium (2 loc./frame), and (c) sparse (1.25 loc./frame) sampling conditions. Dashed vertical lines indicate the mean error for each method. Legend entries show mean and standard deviation. COMET (green, dashed) achieves the narrowest error distribution and lowest mean error across all conditions, with sub-nanometer performance under medium and rich sampling. Methods are: COMET, Minimum Entropy (ME), AIM, RCC (own implementation), RCC (SMAP), and DCC.

#### Supplementary Figure 10

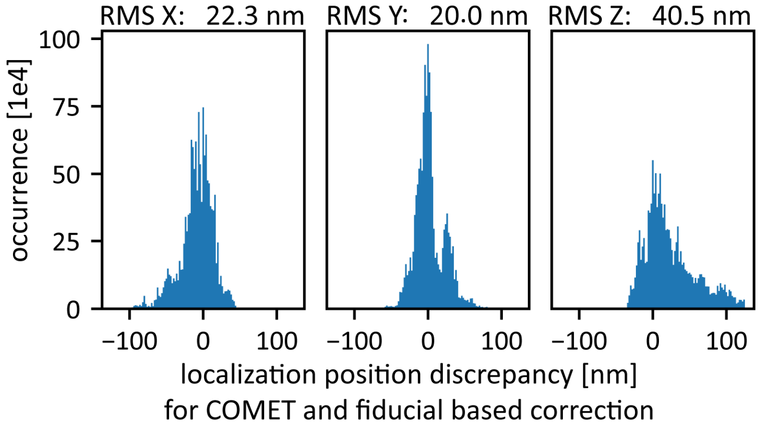

**Supplementary Figure 10: Discrepancy between COMET and fiducial-based corrected localization coordinates in OligoSTORM data.** Histograms of the per-localization position difference between COMET-corrected and fiducial-corrected coordinates in (left) X, (center) Y, and (right) Z. RMS discrepancies are indicated above each panel. The broad distributions, with substantial fractions exceeding 50 nm, reflect systematic differences between the two correction methods, particularly in Z where fiducial tracking is limited by loss of fiducial markers during axial scanning.

Supplementary Figure 11

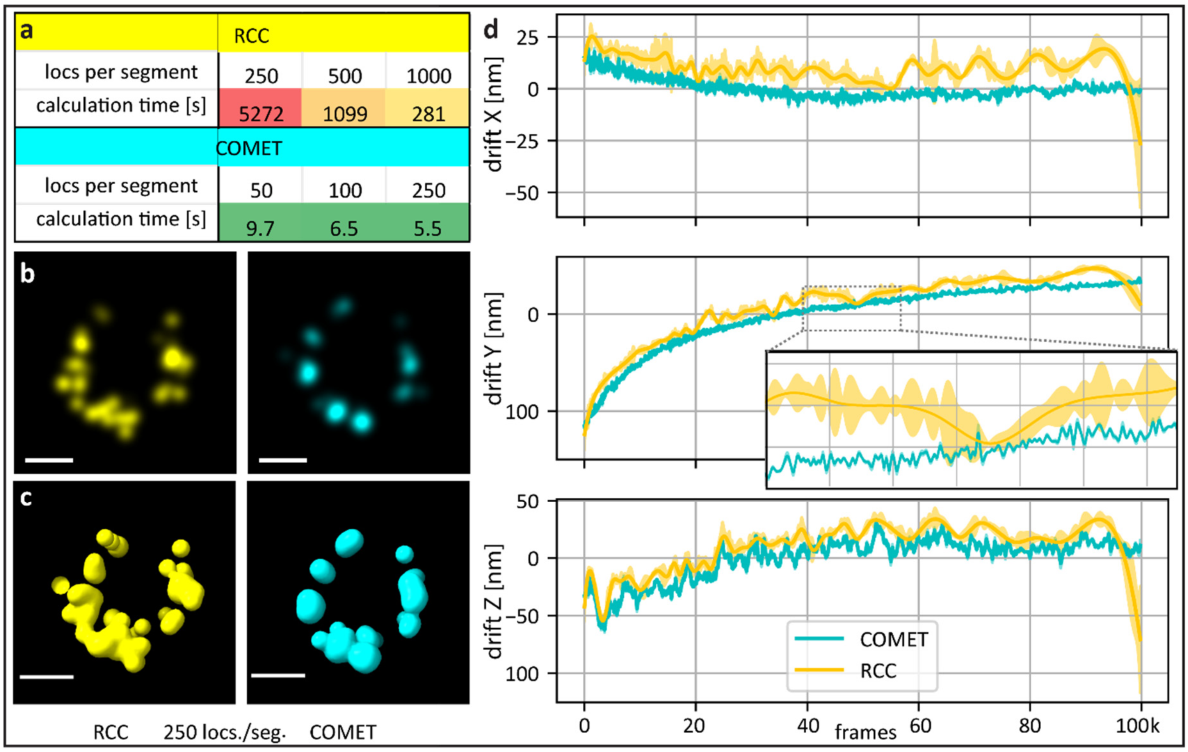

**Supplementary Figure 11: Comparison of COMET and RCC drift correction on experimental 4Pi-STORM data (Nup96).** (a) Computation times for RCC (top) and COMET (bottom) at three segmentation conditions (250, 500, and 1000 localizations per segment). COMET is approximately two orders of magnitude faster. (b) Two-dimensional projections of a single nuclear pore complex unit after drift correction with RCC (left, yellow) and COMET (right, blue) at 250 localizations per time segment. (c) Three-dimensional renderings of the same structure (ChimeraX), showing that the COMET-corrected reconstruction more clearly resolves the expected eightfold symmetry of Nup96. (d) Frame-wise drift estimates from COMET (blue) and RCC (yellow) in X, Y, and Z, with shaded regions indicating the standard deviation across three segmentation conditions. The RCC estimates show substantially larger inter-condition variability, indicating sensitivity to noise in the cross-correlation function. Inset: magnified view of a region where COMET resolves subtle ~10 nm drift transients in Z that RCC does not detect.

#### Supplementary Figure 12

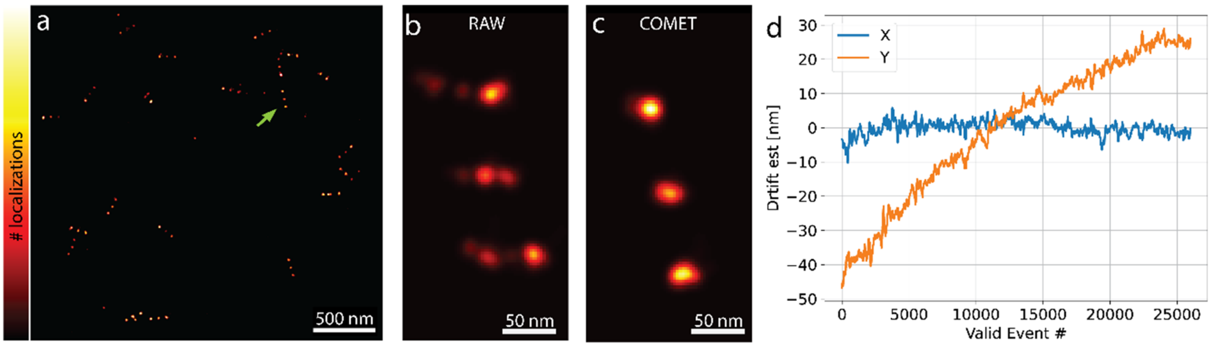

**Supplementary Figure 12: COMET drift correction of MINFLUX nanoruler data.** Overview rendering of Gattaquant 3x80 nm nanorulers acquired using DNA-PAINT MINFLUX, color-coded by localization count. Green arrow indicates the structure shown in panels (b) and (c). (b) Close-up rendering of a single nanoruler before drift correction (RAW), showing drift-induced broadening of the three binding sites. (c) The same structure after COMET drift correction, showing markedly sharper localization clusters consistent with the expected 80 nm inter-site spacing. (d) COMET drift estimate in X (blue) and Y (orange) as a function of valid event number, revealing a continuous linear drift of approximately 25 nm over the course of the measurement despite active stabilization. Scale bars: 500 nm (a), 50 nm (b, c).

#### Supplementary Figure 13

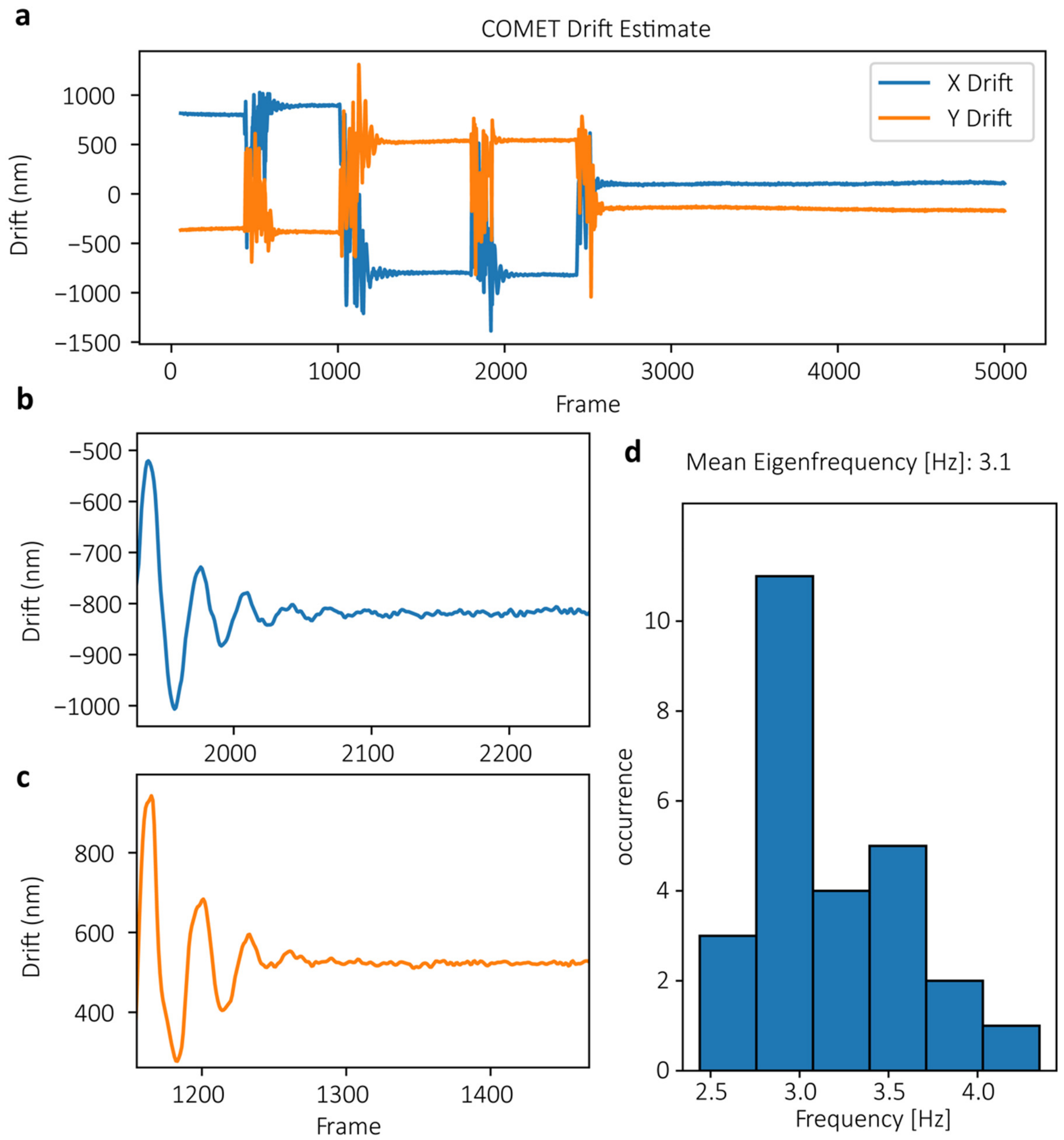

**Supplementary Figure 13: COMET drift correction under deliberately induced extreme mechanical disturbance.** (a) Full drift trajectory estimated by COMET for a 2D STORM dataset acquired during deliberate perturbation of the optical table, showing drift amplitudes exceeding 1 micrometer in both X (blue) and Y (orange). The disturbances consisted of interleaved strong and weak pushes applied during frames ~300--2500. (b, c) Magnified views of the drift estimate immediately following individual disturbance events, revealing damped oscillatory transients with amplitudes of several hundred nanometers and decay times on the order of seconds. (d) Histogram of oscillation frequencies extracted from individual transient events, yielding a mean eigenfrequency of 3.1 Hz, consistent with the mechanical resonance of the floating optical table.

Supplementary Figure 14

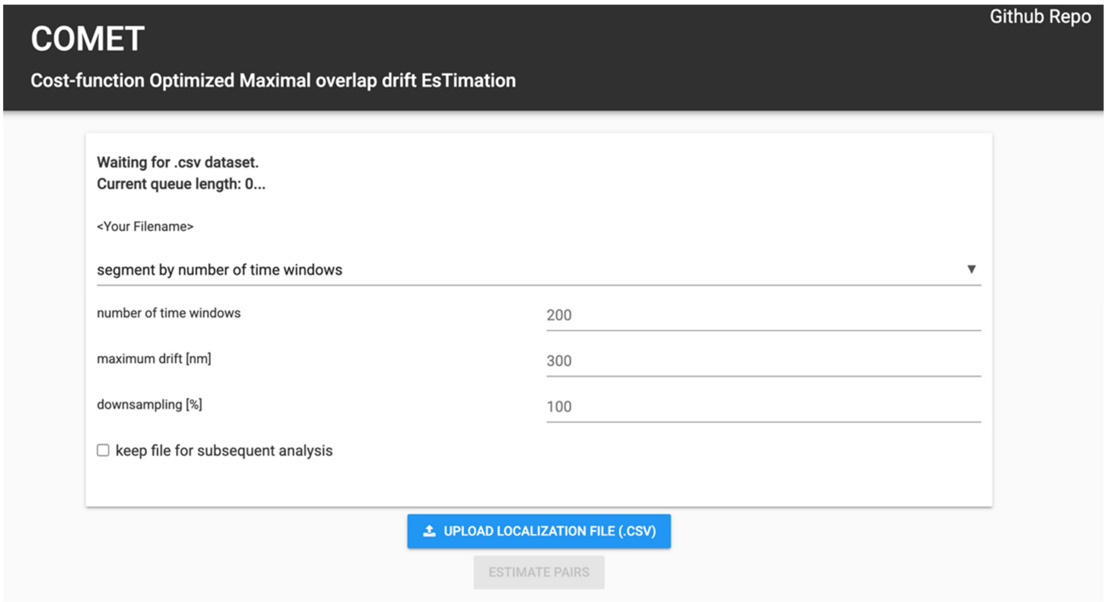

**Supplementary Figure 14: Screenshot of COMET server website.** The web-based COMET service (https://www.smlm.tools) provides a simple, no-installation interface for drift correction of SMLM datasets in standard CSV format.
